# A Temporal Quantitative Profiling of Newly Synthesized Proteins during Aβ Accumulation

**DOI:** 10.1101/2020.01.14.906560

**Authors:** Yuanhui Ma, Daniel B. McClatchy, Salvador Martínez-Bartolomé, Casimir Bamberger, John R. Yates

## Abstract

Accumulation of aggregated amyloid beta (Aβ) in the brain is believed to impair multiple cellular pathways and play a central role in Alzheimer’s disease pathology. But how this process is regulated remains unclear. In theory, measuring protein synthesis is the most direct way to evaluate a cell’s response to stimuli, but to date there have been few reliable methods to do this. To identify the protein regulatory network during the development of Aβ deposition in AD, we applied a new proteomic technique to quantitate newly synthesized protein (NSP) changes in the cerebral cortex and hippocampus of 2, 5 and 9-month-old APP/PS1 AD transgenic mice. This bioorthogonal non-canonical amino acid tagging (BONCAT) analysis combined PALM (Pulse Azidohomoalanine Labeling in Mammals) and HILAQ (Heavy Isotope Labeled AHA Quantitation) to reveal a comprehensive dataset of NSPs prior to and post Aβ deposition, including the identification of proteins not previously associated with AD, and demonstrated that the pattern of differentially expressed NSPs is age-dependent. We also found dysregulated vesicle transportation networks including endosomal subunits, coat protein complex I (COPI) and mitochondrial respiratory chain throughout all timepoints and two brain regions. These results point to a pathological dysregulation of vesicle transportation that occurs prior to Aβ accumulation and the onset of AD symptoms, which may progressively impact the entire protein network and thereby drive neurodegeneration. This study illustrates key pathway regulation responses to the development of AD pathogenesis by directly measuring the changes of protein synthesis and provides unique insights into the mechanisms that underlie AD.

## INTRODUCTION

Alzheimer’s disease is the one of the most common neurodegenerative diseases, and is characterized clinically by a progressive decline in cognitive function and neuropathologically by the presence of neuropil threads, specific neuron loss and synapse loss^1^. Pathological hallmarks include the flame-shaped neurofibrillary tangles of the microtubule binding protein tau and the extracellular plaque deposits of the β-amyloid peptide (Aβ). The detailed pathogenesis of AD is still unclear, primarily due to the complexity of the brain and the challenges of non-invasive studies of pre- and post-symptomatic patients. There are many still postulated or evidenced hypotheses describing the pathophysiology of AD, but efforts to understand AD etiology is complicated by the fact that molecular perturbations vary in different stages of AD and in different brain regions. Short-term memory begins to decline and the cells in the hippocampus degenerate in the early stage of AD while cognitive processes implicated in the cerebral cortex are more gradually altered^2^. Numerous AD studies have determined that Aβ peptide plays a central role in the onset and progression of AD^3^. How Aβ peptides impair cellular pathways and how the brain proteome is remodeled in different brain regions remain unclear.

Analysis of the transcriptome has been widely used to measure a cell’s response to stimuli^4,5^. However, mRNA expression data is not generally consistent with proteome data due to an independent layer of translational regulation^6^. Quantitative proteomic analyses are typically focused on measuring the total quantity of proteins but cannot distinguish the contributions of protein synthesis and degradation to protein levels. Since newly synthesized protein (NSPs) are, in theory, the first to respond to cellular perturbations, measurement of NSPs should permit a more sensitive quantitation of changes than quantitation of the whole proteome. Azidohomoalanine (AHA), a bioorthogonal homolog of methionine, can be metabolically incorporated into proteins by the cell’s own translational machinery. Enrichment of AHA labeled proteins/peptides enable the separation of NSPs from an existing protein pool through a click reaction^7^ with a biotin-linked alkyne followed by enrichment on avidin. Using this AHA labeling strategy, it is possible to improve the temporal resolution of quantitative proteomics. In a previous study, we developed a quantitative strategy using for pulsed azidohomoalanine labeling in mammals (PALM) and applied it to a knock-out mouse model. In PALM, mice were fed with a customized AHA diet within a discrete time period to label NSPs after perturbation^8^. We have also introduced a heavy isotope labeled AHA (H-AHA) to develop the metabolically labeled AHA quantitative strategy HILAQ (Heavy Isotope Labeled AHA Quantitation)^9^. By combining PALM and HILAQ, we were able to achieve efficient and sensitive quantitative labeling in mice through a 4-day L-AHA/H-AHA diet^10^.

To better understand the effects of Aβ peptides on the brain proteome, we applied PALM with H-AHA to the hAPP/PS1 AD mouse model at different ages (2, 5, 9 months). Transgenic hAPP/PS1 mice develop both amyloid deposition and progressive memory loss with age^11^. Here, we performed quantitative NSPs analysis of extracts from hippocampus and cerebral cortex of hAPP/PS1 mice before plaques are formed, after plaques are formed and after the deposition of heavy plaques to find early and potential reversible alterations. We report a novel resource: the newly synthesized proteome of mouse model brain extracts at different stages of AD from 2, 5, and 9-month-old mice. Our quantitative analysis indicated that NSPs are differentially expressed in an age-dependent manner, correlating with the progress of AD pathogenesis. We further found that the dysregulation of endosomal proteins may play an important role in AD pathogenesis and that protein synthesis is not consistent with protein abundance, suggesting that both synthesis and degradation contributed to the protein pool. This study illustrates that quantitation of the newly synthesized proteome is a more direct way to reflect regulation than traditional strategies that focus on protein abundance, and we identify key pathway regulation responses to the development of AD pathogenesis by directly measuring the changes of protein synthesis. Overall, this analysis provides unique insights into the mechanisms that underlie proteomic changes in AD.

## MATERIALS AND METHODS

### Animals/Tissue collection

Three male non-Tg mice and three male borchelt mice (B6.C3-Tg (APPswe,PSEN1dE9)85Dbo/Mmjax) were purchased from the Mutant Mouse Regional Resource Center. All animals were housed in plastic cages located in a temperature- and humidity-controlled animal colony and were maintained on a reversed day/night cycle (lights on from 7:00 P.M. to 7:00 A.M.). Animal facilities were AAALAC-approved, and protocols were in accordance with the IACUC. Harlan laboratories (Envigo) prepared H-AHA and L-AHA pellets. As previously described ^8^, two grams of H-AHA or L-AHA (Cambridge isotope laboratories, Tewksbury, MA) was used to make 1 kg of mouse pellet. Tg-AD mice that were 2, 5, 9 months old and age matched non-Tg mice were given either the H-AHA diet or L-AHA diet for 4 days. The mice were examined daily for gross changes in behavior or physical appearance. All mice were sacrificed by isoflurane inhalation on Day 4. The cerebral cortex (CTX) and hippocampus (HP) were quickly dissected and snap-frozen in liquid nitrogen.

### Tissue Homogenization

The cerebral cortex (CTX) and hippocampus (HP) were transferred to Precellys CK14 tubes containing 1.4 mm ceramic beads and 0.5mm disruption beads (Research Products International Corp.), then 1ml of PBS with protease and phosphatase inhibitors (Roche, Indianapolis, Indiana) was added to CTX and 200 uL was added to HP. The homogenization was performed on Bertin Precellys bead-beating system: 6500 rpm, 3 times, 20 seconds internal. Finally, 0.5% SDS was added and the homogenates were sonicated for 30 secs. A BCA protein assay was performed on each sample.

### Click chemistry

For CTX samples, 2.5mg protein of CTX from H-AHA labeled Tg-AD mice or Non-Tg mice and 2.5mg protein of CTX from L-AHA labeled Non-Tg mice or Non-Tg mice were mixed together as one biological replicate.

For HP samples, 2.5mg protein of HP from H-AHA labeled Tg-AD mice or Non-Tg mice and 2.5mg protein of HP from L-AHA labeled Non-Tg mice or Non-Tg mice were mixed together as one biological replicate.

For each biological replicate, the H-AHA/L-AHA mixture was divided into aliquots (0.25 mg/aliquot). A click reaction was performed on each aliquot as previously published ^12^. Briefly, for each click reaction the following reagents were added in order: 1) 30 ul of 1.7mM TBTA, 2) 8 ul of 50mM Copper Sulfate, 3) 8 ul of 5mM Biotin-Alkyne (C_21_H_35_N_3_O_6_S, Click Chemistry Tools), and 4) 8 ul of 50mM TCEP. PBS was then added to a final volume of 400 ul and reactions were incubated for 1 hour at room temperature. Methanol/Chloroform precipitation was performed, and the precipitated proteins were combined so that there would only be one pellet per 5mg starting material of CTX or 2.5mg of HP.

### Digestion and biotin peptide enrichment

Biotin peptide enrichment was performed as previously described ^13^. Briefly, precipitated pellets were resuspended in 100ul 8M urea and 100ul 0.2% MS-compatible surfactant ProteaseMAX (Promega) in 50mM ammonium bicarbonate, then proteins were reduced, alkylated, and digested with trypsin as previously described ^13^. The digestion was then centrifuged at 13,000 x g for 10 min. The supernatant was transferred to a new tube, and the pellet was resuspended with PBS and centrifuged at 13,000 x g for 10 min. Supernatants were combined, and 150 ul of neutravidin agarose resin (Thermo Scientific) was added. The resin was incubated with the peptides for 2 hours at room temperature while rotating. The supernatant was removed after the resin was spun down *at* 500g for 2 min. The resin was then washed sequentially with 1ml PBS, PBS with 5% acetonitrile, PBS, and distilled water, with a 2 minute spin at 500g after each wash. Supernatants were discarded. Peptides were eluted from the resin twice with 150ul 80% acetonitrile, 0.2% formic acid, and 0.1% TFA on shaker for 5 min at room temperature, and another two times on a shaker at 70 ℃. The resin was spun down at 500g for 2 min between elutions. All elutions were transferred to a single new tube. Prior to MS analysis, the samples were dried in a speed-vac, and dried peptides were resolubilized in buffer A (5% ACN, 95% water, 0.1% formic acid).

### Mass spectrometry

Soluble peptides were pressure-loaded onto a 250-μm i.d. capillary with a kasil frit which had been packed with 2.5 cm of 5-μm Partisphere strong cation exchanger (Whatman) followed by 2.5 cm of 10-μm Jupiter C18-A material (Phenomenex) After loading, the column was washed with buffer A. A 100-μm i.d. capillary with a 5-μm pulled tip packed with 15 cm of 4-μm Jupiter C18 material (Phenomenex) was attached to the loading column with a union, and the entire split-column (loading column–union–analytical column) was placed in line with an Agilent 1100 quaternary HPLC (Palo Alto). The sample was analyzed using MudPIT, using a modified 11-step separation described previously ^14^. The buffer solutions used were: buffer A, buffer B (80% acetonitrile/0.1% formic acid), and buffer C (500-mM ammonium acetate/5% acetonitrile/0.1% formic acid). Step 1 consisted of a 10 min gradient from 0-10% buffer B, a 50 min gradient from 10-50% buffer B, a 10 min gradient from 50-100% buffer B, and 20 min from 100% buffer A. Steps 2 consisted of 1 min of 100% buffer A, 4 min of 20% buffer C, a 5-min gradient from 0-10% buffer B, an 80-min gradient from 10-45% buffer B, a 10-min gradient from 45-100% buffer B, and 10 min of 100% buffer A. Steps 3-9 had the following profile: 1 min of 100% buffer A, 4 min of X% buffer C, a 5-min gradient from 0-15% buffer B, a 90-min gradient from 15-45% buffer B, and 10 min of 100% buffer A. The buffer C percentages (X) were 30, 40, 50, 60, 70, 80, and 100%, respectively, for the following 7-step analysis. In the final two steps, the gradient contained: 1 min of 100% buffer A, 4 min of 90% buffer C plus 10% B, a 5-min gradient from 0-10% buffer B, an 80-min gradient from 10-45% buffer B, a 10-min gradient from 45-100% buffer B, and 10 min of 100% buffer A. As peptides eluted from the microcapillary column, they were electrosprayed directly into an Orbitrap Elite mass spectrometer (Thermo Scientific) with the application of a distal 2.4-kV spray voltage. A cycle of one full-scan FT mass spectrum (300-1600 m/z) at 240,000 resolution, followed by 20 data-dependent IT MS/MS spectra at a 35% normalized collision energy, was repeated continuously throughout each step of the multidimensional separation. Application of mass spectrometer scan functions and HPLC solvent gradients were controlled by the Xcalibur data system.

### Data analysis

Both MS1 and MS2 (tandem mass spectra) were extracted from the XCalibur data system format (.RAW) into MS1 and MS2 formats using in-house software (RAW_Xtractor)^15^. MS/MS spectra remaining after filtering were searched with ProLuCID ^16^ against the UniProt_Mouse_03-25-2014 database concatenated to a decoy database, in which the sequence for each entry in the original database was reversed ^17^. All searches were parallelized and performed on a Beowulf computer cluster consisting of 100 1.2-GHz Athlon CPUs ^18^. No enzyme specificity was considered for any search. The following modifications were searched for analysis: a static modification of 57.02146 on cysteine for all analyses, a differential modification of 452.2376 on methionine for AHA-biotin-alkyne, or 458.2452 for H-AHA-biotin-alkyne. ProLuCID results were assembled and filtered using the DTASelect (version 2.0) program ^19,20^. DTASelect 2.0 uses linear discriminant analysis to dynamically set XCorr and DeltaCN thresholds for the entire dataset to achieve a user-specified false discovery rate (FDR). In DTASelect, the modified peptides were required to be fully tryptic, less than 5-ppm deviation from peptide match, and identified with an FDR at the spectra level of 0.01. The FDRs are estimated by the program from the number and quality of spectral matches to the decoy database. For all datasets, the protein FDR was < 1% and the peptide FDR was < 0.5%. The MS data was quantified (i.e., generate heavy/light ratios) using the software pQuant^21^, which uses the DTASelect and MS1 files as the input. pQuant assigns a confidence score called sigma to each heavy/light ratio from zero to one. zero, the highest confidence, means there is no interference signal, whereas one means the peptide signals are almost inundated by interference signals (i.e., very noisy). For this analysis, only peptide ratios with sigma less than 0.5 were used for quantification.

### Pathway analysis

Functional pathway analysis and biological processes enrichment were performed using the online tool STRING (https://string-db.org/cgi/network.pl). Protein networks from STRING were visualized by Cytoscape. Colors were defined to show fold of change. Shapes were defined to indicate timepoints and brain regions.

### Western blots

For total homogenates: Protein were resolved by NuPAGE 4-12% Bis-Tris gel (Life technologies) and transferred onto PVDF membranes using a semi-dry blotting apparatus (Life technologies). After blocking with 5% milk powder in PBS with 0.05% Tween 20, the membranes were incubated overnight at 4 °C, first with primary antibodies and then with an HRP-conjugated secondary antibody.

For confirmation of protein levels in NSP datasets: 5mg proteins were taken as starting material for each sample, followed by click reaction and Methanol/Chloroform precipitation. Precipitated pellets were resuspended in 500ul 8M urea. After centrifugation, the supernatants were transferred to a new tube and the pellets were dissolved in 5% SDS followed by addition of 400ul PBS. The supernatants were combined and incubated with 80ul neutravidin beads for two hours at room temperature while rotating. The resin was then washed sequentially with 1-ml PBS, PBS with 5% acetonitrile, PBS, and a final wash of distilled water. The resin was spun down at 500g for 2 min after each wash. The proteins were eluted from the resin by boiling in 50ul 4×sample buffer (Bio-Rad) and 2.5ul 20×reducing reagent (Bio-Rad). Protein were resolved by 4-12% gradient SDS-PAGE and transferred onto PVDF membranes using a semi-dry blotting apparatus (Life technologies). After blocking with 5% milk powder in PBS with 0.05% Tween 20, the membranes were incubated overnight at 4 °C with primary antibodies and then with an HRP-conjugated secondary antibody. Protein bands were visualized by chemiluminescence. The following primary antibodies were used in this study: mouse monoclonal anti-Aβ1-16 (803001; Biolegend), monoclonal mouse anti-UBE3A (sc-166689; Santa Cruz), mouse monoclonal anti-EHD3 (sc-390513; Santa Cruz), mouse monoclonal anti-CMPK1 (sc-376153; Santa Cruz) The following secondary antibodies were used in this study: HRP conjugated goat anti mouse secondary antibody (ab97023; Abcam).

## RESULTS

### Confirmation of AD pathogenesis

The hAPP/PS1 AD mice and non-transgenic wild type littermates were fed a diet of either light AHA (L-AHA) or heavy AHA (H-AHA) for four days at 2, 5 and 9 months. The cerebral cortex and hippocampus were quickly dissected at day 4 of labeling, rinsed with PBS, and snap frozen. Each brain region was separately homogenized (Figure 1A). Before proteomic analysis, AD pathogenesis at the three different time points was confirmed in APP/PS1 mice. ELISA was performed to measure the amount of Aβ in both cerebral cortex and hippocampus. Data shown in Figure 1B reveals that there were no significant differences in Aβ levels in cerebral cortex and hippocampus at 2 months. At 5 months and 9 months, Aβ levels were increased with the age of APP/PS1 mice in both brain regions compared to the non-transgenic littermates. These results are consistent with previous reports, which show that Aβ deposition starts to form at 5-6 months in this model^22^. Moreover, Aβ levels in two brain regions did not show significant differences at 5 months while Aβ levels in cerebral cortex were slightly higher than that in hippocampus in APP/PS1 mice at 9 months, suggesting different physiopathological activities. To measure AHA incorporation in mice at different time points, we mixed equal amounts of proteins from H-AHA and L-AHA labeled mice and performed dot blots. We observed no significant difference in total protein synthesis between cerebral cortex and hippocampus, but we saw a decline in AHA incorporation in both cerebral cortex and hippocampus with time (Figure 1D and 1E). Overall, these experiments confirm AHA incorporation in this disease model and verify Aβ deposition in both cerebral cortex and hippocampus. Based on the measurement of Aβ accumulation, previous pathological and behavioral analyses^22^, we considered 2 months, 5months and 9 months to be the best time points to represent early-, mid-, and late-stages of AD.

**Figure 1.**
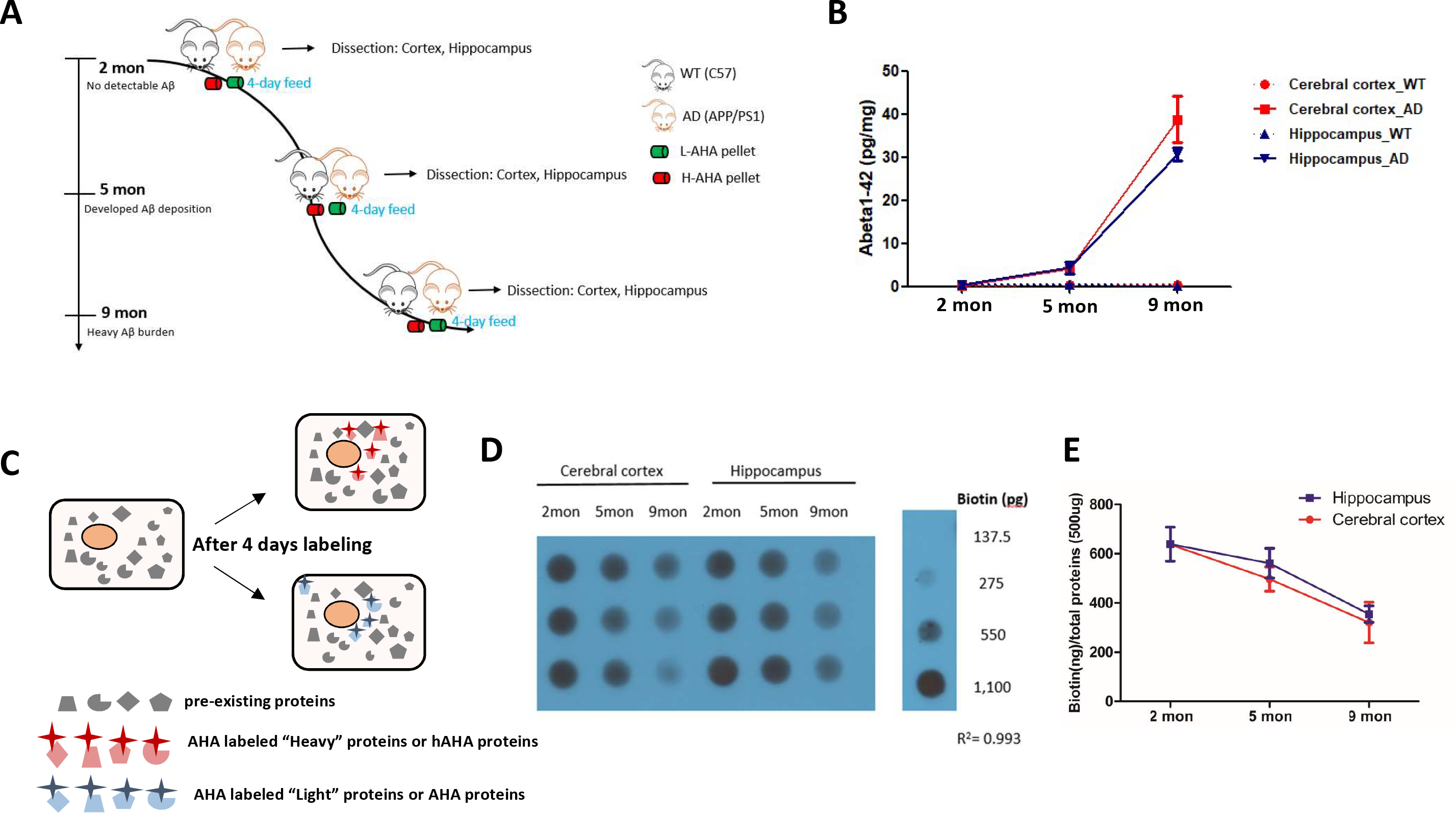
Schematic of labeling and AD pathogenesis confirmation. **A**: Pulse labeling of AHA/hAHA in mice. After feeding mice with AHA or hAHA diet for 4 days at 2, 5, 9 months, the mice were sacrificed and the tissues were extracted. **B**: ELISA analysis confirmed that Aβ levels are increased in both cerebral cortex and hippocampus along with the age of APP/PS1 mice. The cortex and hippocampus homogenates were assayed at 2, 5 and 9 months for human-specific Aβ42.Bars, means±S.D. (n=3). **C**: The schematic diagram of the AHA/hAHA labeling in mice cells through 4 days of feeding. **D**: Incorporation of AHA/hAHA is decreased with age in both cerebral cortex and hippocampus. Equal amount of proteins from hAHA and AHA labeled mice were mixed. Biotin can be added to AHA/hAHA proteins through click reaction and its detection can reflect the Incorporation of AHA/hAHA in cerebral cortex and hippocampus of 2, 5, 9 months mice by dot blot. **E**: The grayscale of dots in **D** were measured by image J and was presented by the points. Bars, means±S.D. (n=3).

Next, L-AHA and H-AHA tissues were mixed 1:1 (wt/wt). APP/PS1 AD mouse cerebral cortex or hippocampus was mixed with an age-matched control. Three biological replicates were analyzed for each L-AHA/H-AHA mixture (six mice per time point). For each time point, a label swap was performed between conditions to check for any possible effects of the heavy label on the proteome. Click chemistry was performed on these mixtures to covalently add a biotin-alkyne. After tryptic digestion, the AHA peptides were isolated with neutravidin beads. Modified peptides were eluted from the beads and then identified by MS.

### Identification of newly-synthesized proteins in both cerebral cortex and hippocampus at 2, 5, 9 months and overall similarity

Consistent with our dot-blot results, a similar decrease in the number of identified NSPs with age in both cerebral cortex and hippocampus was observed, suggesting that AHA incorporation through feeding had been reduced with age (Figure 2A, **supplementary Table 1**). Additionally, there is no significant difference in the number of NSPs between AD and WT for each timepoint. (Figure 2B). Next, ion chromatograms for the light and heavy AHA peptide pairs were extracted, and heavy/light ratios were calculated. In each group, 85-93%% of identified NSPs were quantified with normal distributions (**Supplementary Figure 1B and supplementary Table 2**). NSPs quantified in all three replicates were considered for further analysis (Table 1). There were 247 and 210 NSPs quantified in all biological replicates of all time points for cerebral cortex and hippocampus, respectively (Figure 2C&D). With the global map of NSPs in hand we first performed a hierarchical clustering of the proteins quantified in all time points. We observed that the 9 months sample exhibited a strong divergence from samples at the 2 and 5 month time points in both cerebral cortex and hippocampus (Figure 3A&B). The heat-map in Figure 3A&B shows that the difference is driven heavily by protein clusters 3 and 6 in both brain regions.

**Table 1.** Number of quantified NSPs in Cerebral cortex and Hippocampus. The group at each time point were consisted of 3 biological replicates, in which there are a pair of label swap treatments and one forward or reverse treatment. Proteins quantified in all three replicates are considered to be reliable for further analysis.

| Timepoint | Biological Replicates | Content | Proteins quantified in all three biological replicates |  |
| --- | --- | --- | --- | --- |
|  |  |  | Cerebral Cortex | Hippocampus |
| 2mon | 3 | H-AD/L-WT<br>L-AD/H-WT<br>L-AD/H-WT | 1,549 | 1,156 |
| 5mon | 3 | H-AD/L-WT<br>L-AD/H-WT<br>L-AD/H-WT | 908 | 523 |
| 9mon | 3 | L-AD/H-WT<br>H-AD/L-WT<br>H-AD/L-WT | 337 | 377 |

**Figure 2.**
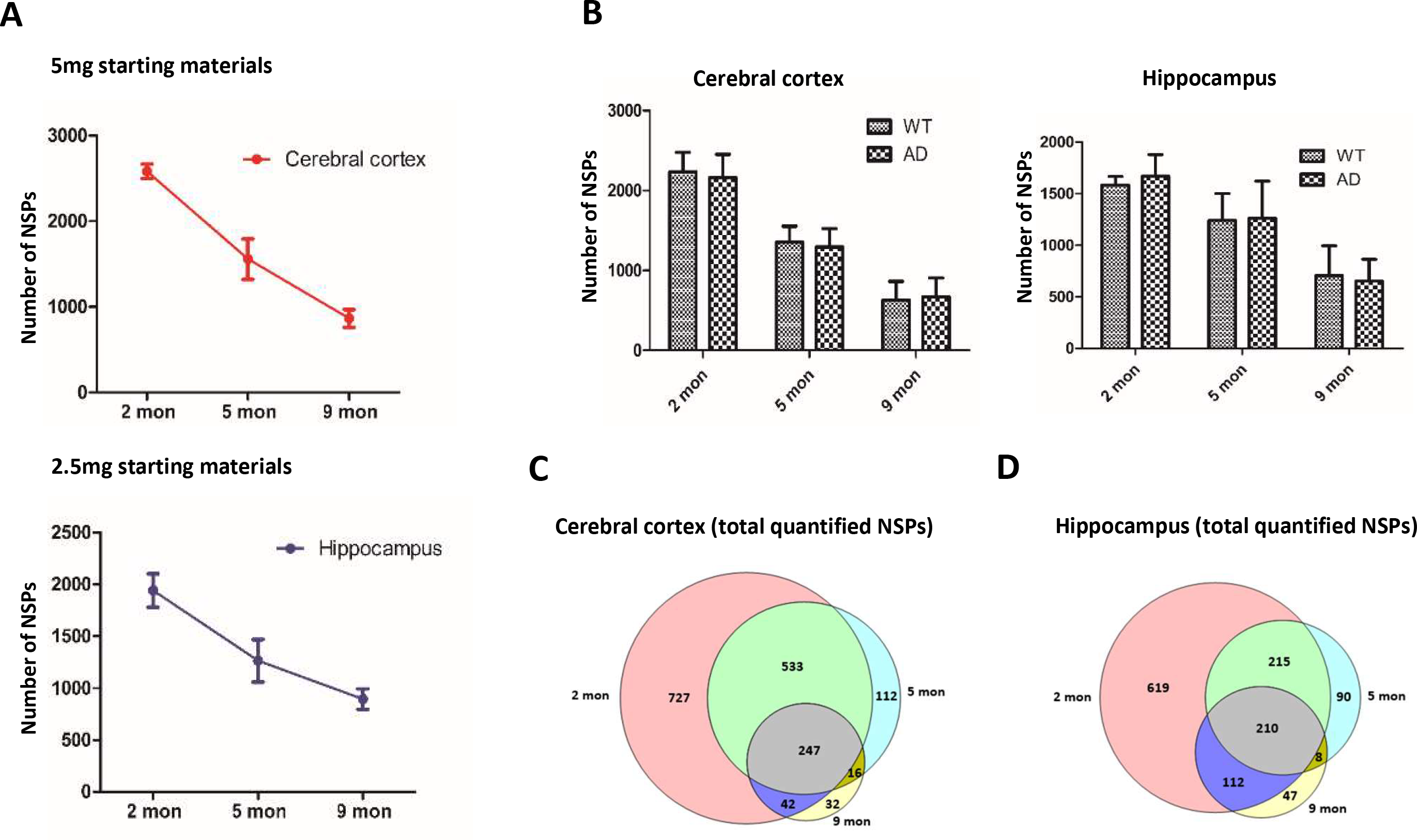
Identification and quantitative results **A**: The number of NSPs identified from an equal amount of AHA /hAHA protein mix by mass spectrometer are decreased with age in both cerebral cortex and hippocampus. Bars, means±S.D. (n=3). **B**: There is no significant difference in the number of NSPs between AD and WT in different age of mice. Bars, means±S.D. (n=3). C&D. Venn diagram analysis for total quantified proteins at 2, 5, 9 months in cerebral cortex and hippocampus.

**Figure 3.**
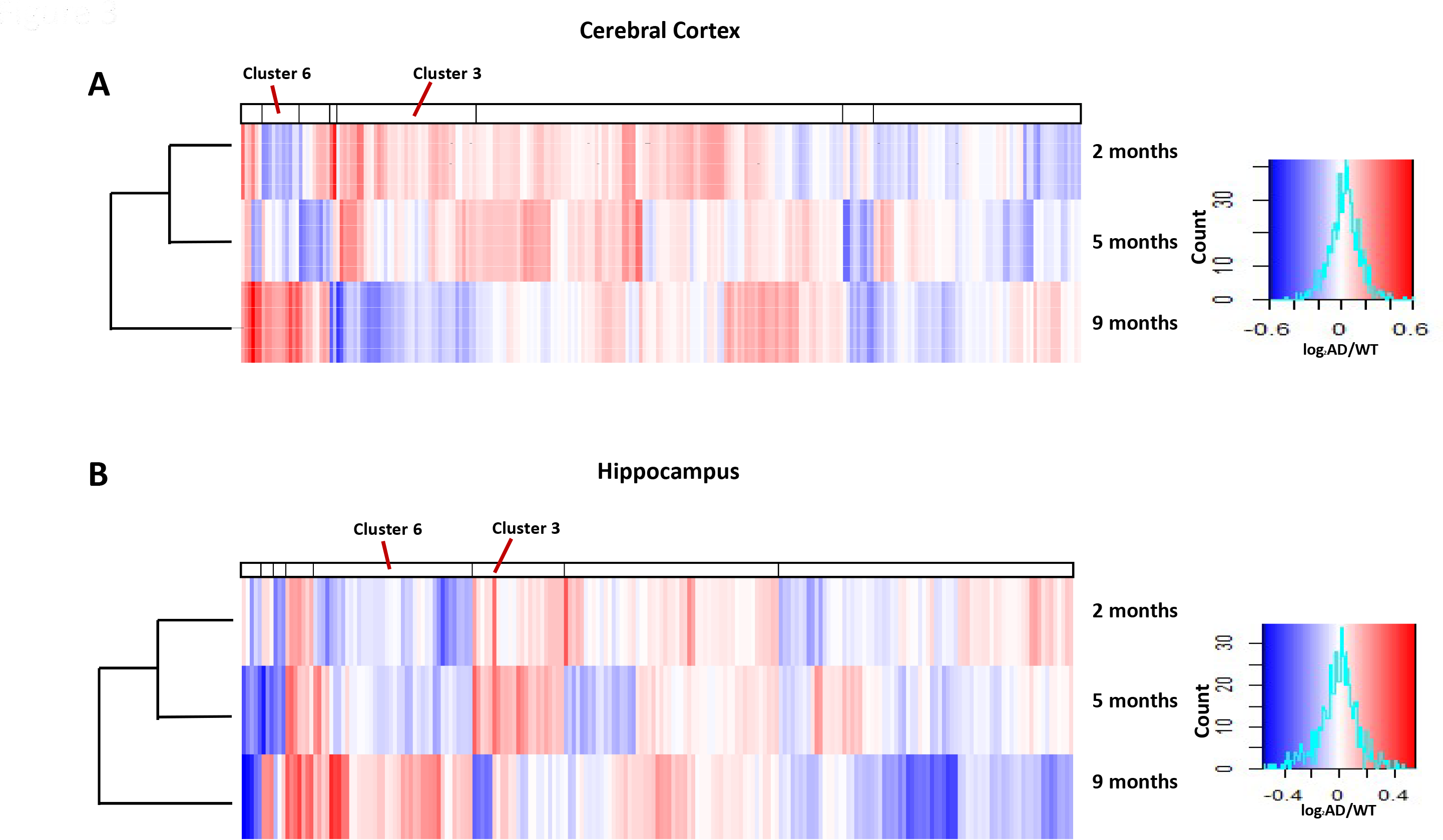
Cluster analysis. Heat map showing mean-normalized protein fold changes for 247 overlapped proteins and 210 proteins over all time points in cerebral cortex (**A**) and hippocampus (**B**), respectively.

For cerebral cortex, enrichment analysis on cluster 3 showed that the most significantly associated biological process is related to synaptic vesicle cycle (**Supplementary Figure 2 and Supplementary Table 3**), including excitatory amino acid transporter 1 (GluT1), excitatory amino acid transporter 2 (GLT1), synaptosomal associated protein 25 (SNAP25), AP2 complex subunit beta (AP100B) and AP2 complex subunit alpha-1 (AP100A). These five NSPs were all down-regulated at 9 months. Lack of GLT1, which is a widely distributed astrocytic glutamate transporter, can result in epilepsy and exacerbation of brain injury ^23^. Recent studies reported that increased GLT1 expression in activated astrocytes may exert beneficial roles in attenuating inflammation-induced memory loss and preserving cognitive function, even in presence of Aβ ^24^. SNAP25 had been proposed to be a promising biomarker for synapse degeneration in AD^25^. As a cargo receptor, AP2 plays an important role in endocytosis activity. These data suggested attenuated synaptic vesicle transportation at 9 months, which may play a crucial rule in inducing severe AD pathology. Enrichment analysis on cluster 6 showed increased expression of NSPs associated with the process of Fatty acid metabolism and RNA transport at 9 months (**Supplementary Figure 3 and Supplementary Table 3**). These expression differences supported previous findings suggesting that fatty acid metabolism is significantly dysregulated in AD brains^26^. For hippocampus, the most significant biological processes enriched for NSPs in cluster 3 are mainly bioenergetics, including glycolysis, gap junctions, carbon metabolism and pentose phosphate pathway (**Supplementary Figure 4 and Supplementary Table 4**). Lower levels ofglycolysis (cluster gene: Gpi, Aldoc, ldhb) at 9 months compared with earlier time points is consistent with the previous study, which claimed lower rates of glycolysis and higher brain glucose levels correlated to more severe plaques and tangles found in the late-onset AD brains ^27^. In protein enrichment analysis of cluster 6, we noticed that one of the most significant pathways referred to AD (**Supplementary Figure 5 and Supplementary Table 4**), in which prolow-density lipoprotein receptor-related protein 1 (LRP1) and Apoliprotein E (ApoE) were enriched. APOE isoforms are genetic risk factors for AD and has an important role in Aβ metabolism^28^. Studies show that APOE genotypes strongly affect deposition of Aβ to form senile plaques and cause cerebral amyloid angiopathy, two major hallmarks of amyloid pathology in AD brains ^29^. Over the past decade there has been more focus on the role of LRP1, especially in late-onset AD. LRP1 not only regulates the metabolism of Aβ in the brain and periphery, but also maintains brain homeostasis, impairment of which likely contributes to an Aβ-independent development of AD development. Several preclinical studies have also demonstrated LRP1 involvement in regulating the pathogenic role of APOE.

**Figure 4.**
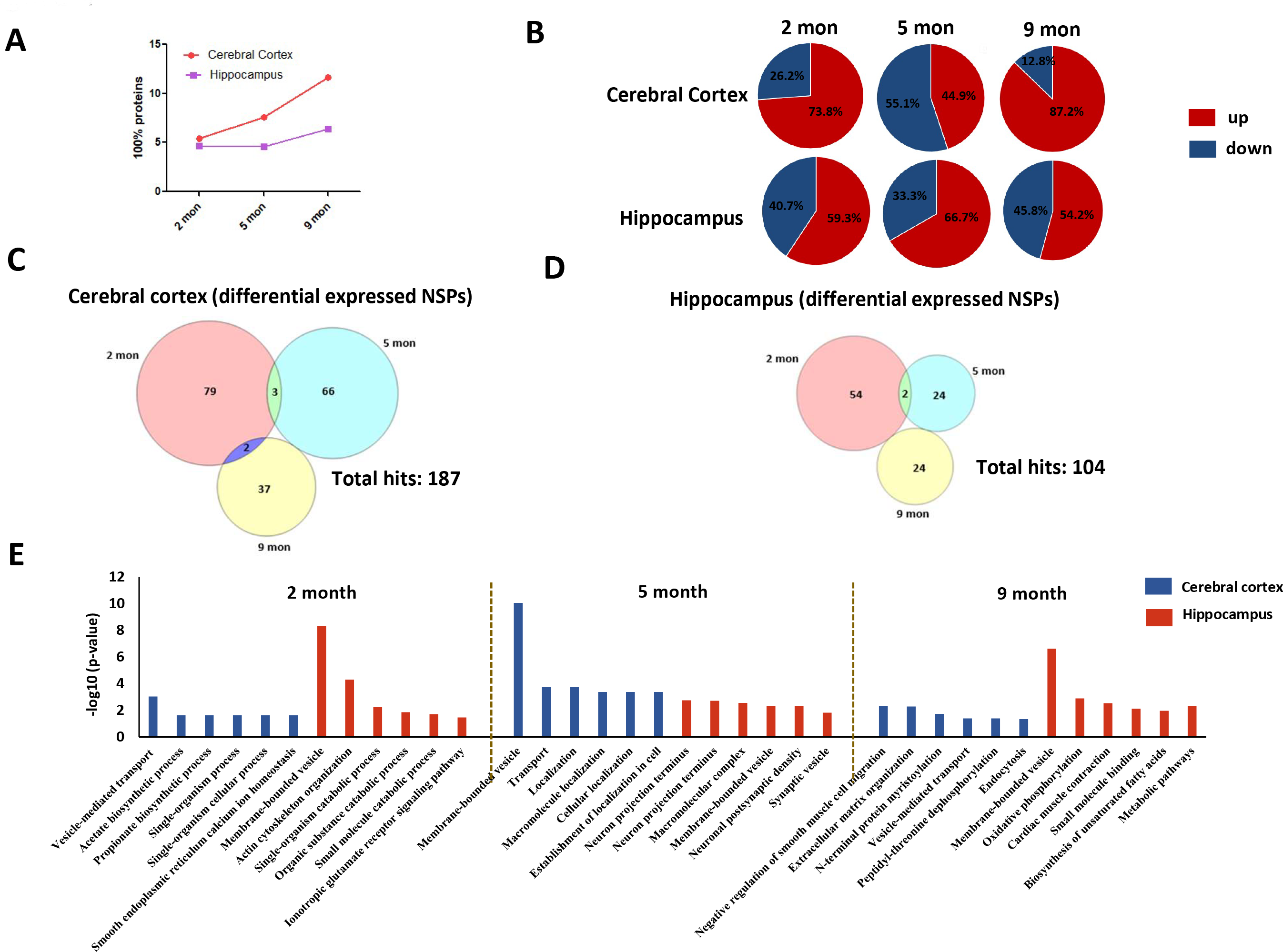
Differentially expressed changes. **A**. Graphs of age-dependent increase of the percentage of significantly altered proteins in total quantified datasets. **B**. The distribution of significantly upregulated and down-regulated proteins at each time point. **C&D**. Venn diagram analysis for significantly changed proteins at 2, 5, 9 months in cerebral cortex and hippocampus. **E**. Biological process annotation on the significantly altered proteins at 2, 5, 9 months in cerebral cortex and hippocampus using STRING database. The 6 biological processes identified with the lowest false discovery rate (FDR) are displayed.

**Figure 5.**
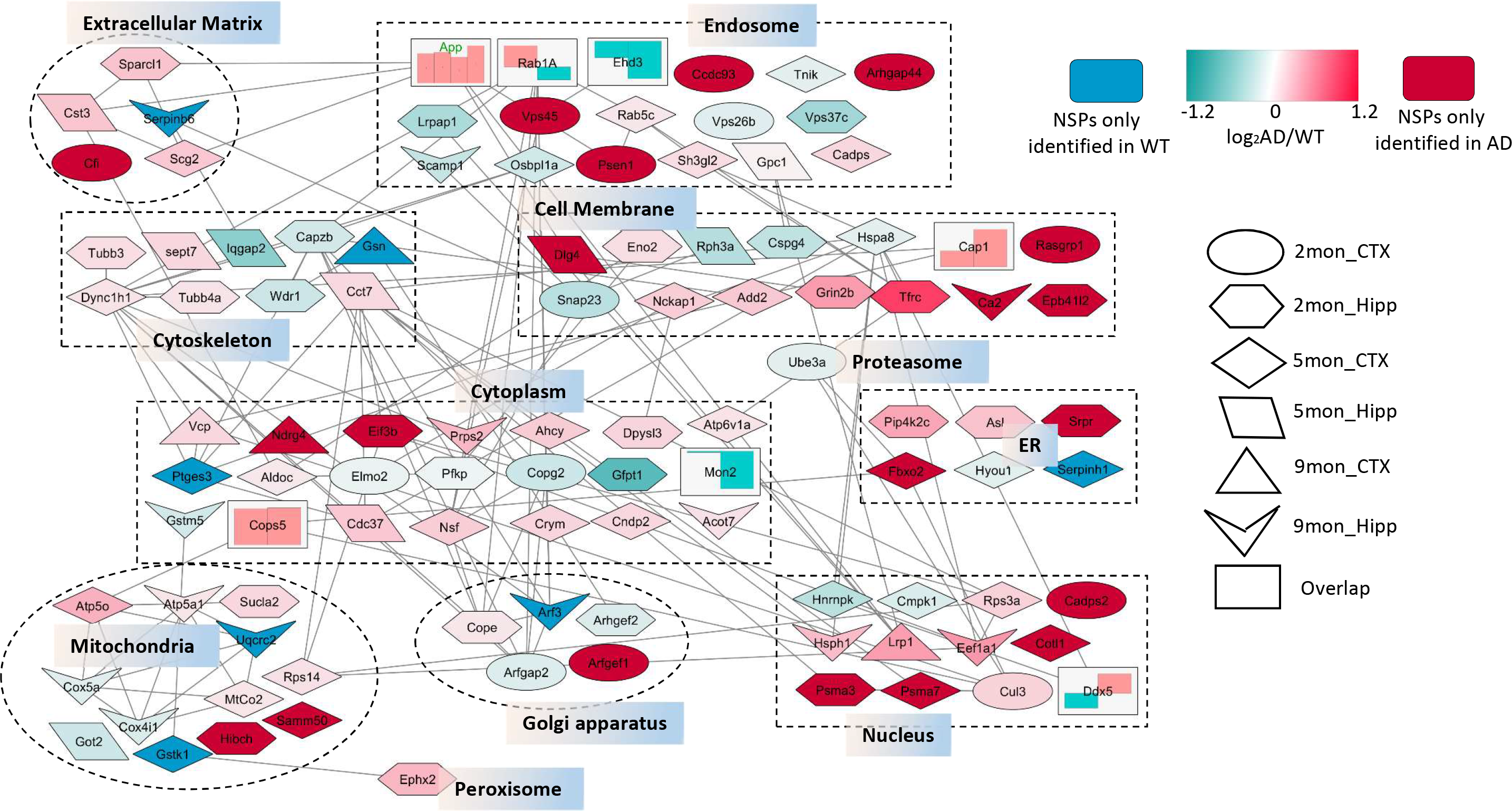
STRING network analysis of significant altered NSPs, which were enriched in vesicle related processes. Only interactions with a STRING score ≥ 0.7 are shown. Node colors are linearly related to fold-change (Rectangular node excluded). Node shapes represent different timepoints and brain region. The shape “rectangular” represents the overlapped NSPs fold-changes indicated in bar graph. The column in red means increased synthesis while that in green means down regulation. The overlapped NSPs were listed here with bar graph annotated from left to right. **APP**: 2mon_CTX; 2mon_Hipp; 5mon_Hipp; 9mon_CTX; **Cap1**: 2mon_Hipp, 9mon_CTX; **Cops5**: 2mon_Hipp, 5mon_Hipp; **Ddx5**: 5mon_CTX; 9mon_Hipp; **Ehd3**: 5mon_CTX; 9mon_Hipp; **Mon2**: 2mon_Hipp; 5mon_CTX; **Rab1A**: 9mon_CTX; 9mon_Hipp

### Age-specific alteration of newly-synthesized proteome in hAPP/PS1 brains

We considered proteins to be changed if they fit into the following categories: (1) proteins which have statistically significant changes, with p<0.05; (2) proteins with average fold change more than 1.5; (3) proteins with reliably unquantified large changes. Due to the limited dynamic range of mass spectrometry, when there are very large changes between a light and heavy protein, only the more abundant is detected and quantitation is impossible. Unquantified proteins that were only identified in one condition and not in the other biological replicates were considered to have reliably large changes^10^. After combining the proteins from all three categories, we found that 5.40%/4.67%, 7.59%/4.60% and 11.6%/6.38% of total protein were changed significantly in 2, 5, 9 month hAPP/PS1 AD cerebral cortexes/hippocampus (Table 2).The percentage of reliable significantly changed proteins in cerebral cortex indicated a sharper increase with age than in hippocampus (Figure 4A). In both cerebral cortex and hippocampus, there were more up-regulated proteins present at almost all time points (Figure 4B). Furthermore, Venn diagram analysis indicated that, in both cerebral cortex and hippocampus, the overlap of differentially expressed NSPs among different timepoints is far less (Figure 4C&4D) than that of total quantified NSPs (Figure 2C&4D), suggesting that the pattern of differentially expressed NSPs is age-dependent and changed as AD pathogenesis progressed.

**Table 2.** The list of differential expressed NSPs. The proteins which has statistical significant changes with p<0.05 or very large changes with average fold change more than 1.5 or unquantified large changes * were counted as reliable changes for the further analysis. P value were calculated by one sample t test.

| Cerebral Cortex |  |  | Age | Hippocampus |  |  |
| --- | --- | --- | --- | --- | --- | --- |
| Total | Up | Down |  | Up | Down | Total |
| 84 | 62 | 22 | 2 mon | 32 | 22 | 54 |
| 69 | 31 | 38 | 5 mon | 16 | 8 | 24 |
| 39 | 34 | 5 | 9 mon | 13 | 11 | 24 |

\*When there are very large changes between a light and heavy protein, only one is detected by the mass spectrometer due to its limited dynamic range.

### Functional network of significant altered NSPs enriched in vesicle transport

Gene ontology (GO) analysis was performed on changed NSPs from above three categories at 2, 5 and 9 months. Most interestingly, for the six analyses of cerebral cortex and hippocampus throughout all time points, vesicle mediated transport/membrane bounded vesicle was included in all the top 6 pathways with significant p values (p<0.05) (Figure 4E). Moreover, the significantly altered NSPs enriched in vesicle mediated transport/membrane bounded vesicle constituted a larger proportion in hippocampus than that in cerebral cortex (**Supplementary Figure 6**). The previous study showed that vesicle transport is involved in APP trafficking network and regulates amyloidogenic processing pathway ^30^. Additionally, vesicle mediated Aβ internalization and degradation in glial cells is one of the most important ways that Aβ levels are regulated in the brain^31,32^

**Figure 6.**
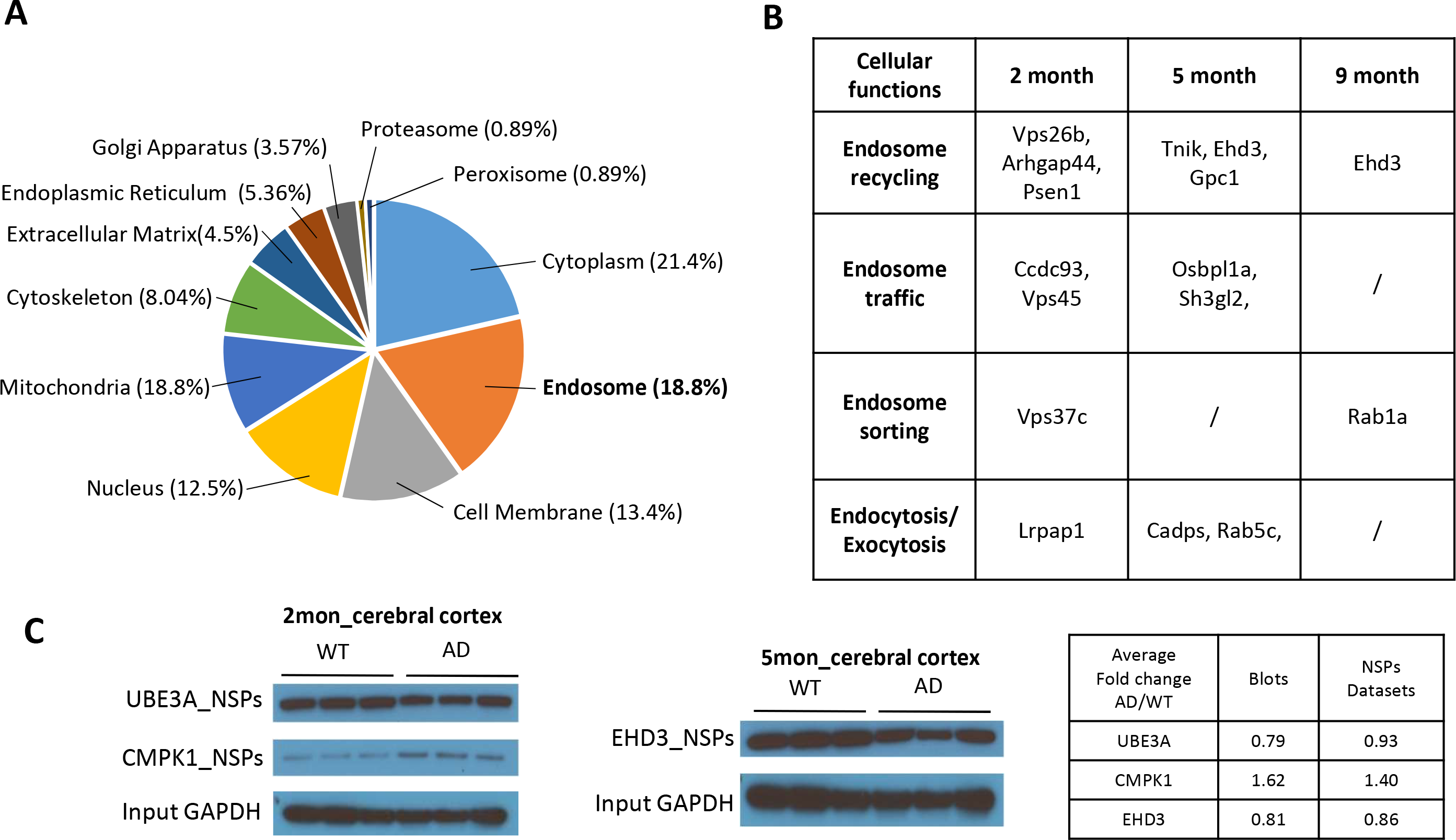
**A:** The subcellular distribution of significantly changed NSPs which were enriched in vesicle transportation process. **B**: More detailed annotation of endosomal proteins from Figure 6A according to previous publications. (**Endosome recycling**: Vps26b^51^, Arhgap44^52^, Psen1^53^, Tnik^54^, Ehd3^55^, Gpc1^56^; **Endosome traffic**: Ccdc93^57^, Vps45^58^, Osbpl1a^59^, Sh3gl2^60^; **Endosome sorting**: Vps37c^61^, Rab1a^62^; **Endocytosis/Exocytosis**: Lrpap1^63^, Cadps^64^, Rab5c^65^. **C**: Verification of NSPs changes by western blot. The proteins of each sample were performed click reaction and Methanol/Chloroform precipitation. Then NSPs were enriched using neutravidin beads and eluted from the resin by boiling in 50ul 4×sample buffer and 2.5ul 20×reducing reagent. Protein were resolved by 4-12% gradient SDS-PAGE and transferred onto PVDF membranes using a semi-dry blotting apparatus. After blocking with 5% milk powder in PBS with 0.05% Tween 20, the membranes were incubated overnight at 4 °C with primary antibodies and then with an HRP-conjugated secondary antibody. Protein bands were visualized by chemiluminescence.

When all timepoints and brain regions are combined, there were in total 105 significantly changed NSPs enriched in vesicle related processes (**Supplementary Table 5**). To further characterize the significant *de novo* proteomic changes in vesicle transport observed in hAPP/PS1 AD mice, we performed a network analysis using STRING on the 105 NSPs (Figure 5). The proteins were annotated with multiple subcellular compartments, including cytoplasm, endosome, cell membrane, nucleus, mitochondria, cytoskeleton, endoplasmic reticulum (ER), extracellular matrix (EM), Golgi apparatus, proteasome and peroxisome (Figure 6A). NSPs from the 2 months timepoints distributed in these subcellular compartments and also interacted with those from later timepoints in combined vesicle transportation network, suggesting that early regulation occurs in multiple loci prior to Aβ deposition and spreads further via protein interactions. We noticed that more than 50% of NSPs in this network were enriched in cytoplasm, endosome and cell membrane. The most striking part of this analysis is the enrichment of endosomal proteins throughout all timepoints and both brain regions. Endosomes are membrane bounded vesicles, which are formed via a complex process known as endocytosis. We further annotated the endosomal proteins into four processes including endosome recycling, endosome traffic, endosome sorting and endocytosis/exocytosis (Figure 6B). Endosome recycling is coordinated with endocytosis/exocytosis to control the abundance and topography of surface receptors^33^. Disruption in the endocytosis process has been implicated in AD pathogenesis^34^. It has been reported that dysfunction of endosomes engenders early signs of neurodegeneration including activation of Ras-related protein 5 (RAB5) isoforms, which causes enlargement of early endosomes and disruption of retrograde axonal trafficking of nerve growth factor (NGF) signals^34^. Vacuolar protein sorting 45 (VPS45), which is recruited to early endosomal membrane by Rab5 isoforms^35^, also plays an important role in endocytosis. The synthesis of both of VPS45 and RAB5C in our dataset were significantly increased at 2months and 5months, which was consistent with the previous reports. In addition, EH domain containing protein 3 (EHD3) can regulate the dendritic transport of β-secretase (BACE1), and its depletion of EHD3 in hippocampal neurons can decrease Aβ production ^36^. We found that EH domain containing protein 3 (EHD3) was synthesized less in both 5 months cortex and 9 months hippocampus of AD mice. This could represent adaptive down-regulation of EHD3 to help reducing the stress from Aβ over-production in the later timepoints.

Finally, we further confirmed that the synthesis of EHD3, UMP-CMP kinase 1 (CMPK1) and ubiquitin-protein ligase E3A (UBE3A) were altered consistently with the proteomic dataset in our study (Figure 6C), indicating the changes detected by MS are reliable. Together, our results demonstrated that vesicle transportation plays a pivotal role in AD because it regulates distinct proteins in the same functional groups as Aβ accumulation and AD pathogenesis.

## Discussion

Aβ deposition elicits a range of toxic effects in mouse brain. To study the progress of alterations that underlie AD pathogenesis caused by Aβ accumulation from such a complex proteome, we have applied PALM combined H-AHA labeling strategy and quantitative mass spectrometry-based proteomic analysis in hAPP/PS1 mouse model in which Aβ was overexpressed. Here, we identified thousands of NSPs in hippocampus and cerebral cortex from 2, 5, 9 months hAPP/PS1 AD mouse and wildtype controls, and we found an age-specific pattern of the significant NSPs alterated in both brain regions. Vesicle transport related processes showed up in each GO analysis of significantly changed NSPs from different timepoints and brain regions. Endosomal proteins constituted a large proportion of the vesicle transport network in our dataset and were mainly enriched in endosome recycling, endosome traffic, endosome sorting and endocytosis/exocytosis. Furthermore, we confirmed the alteration of specific proteins.

To study the progression of alterations that underlie AD pathogenesis, we sought to reduce complexity by quantifying the NSP sub-proteome using the methionine surrogate AHA and its heavy labeled version. AHA has been routinely employed to identify and quantify NSPs by MS in cell culture^9^. Recently several studies have reported the use of AHA labeling in mouse to examine de novo protein synthesis during a discrete window^8,37^. Heavy isotopes such as heavy amino acid^38^ and heavy water^39^ have also been used to track changes in protein synthesis. Our approach has several advantages over previous methods. Labeled NSPs can be purified from the whole proteome through biotin enrichment, thereby increasing the detection of low abundant NSPs. In other methods, the abundance of unlabeled “old” proteins will reduce NSPs detection by MS because of its limited dynamic range. Furthermore, we introduced heavy isotope labeled AHA in mice, which enables NSPs enrichment, confirmation, and quantification using a single stable isotope-labeled molecule.

With our strategy, NSPs changes in response to Aβ accumulation were quantified in age-different hAPP/PS1 AD mice with paired control. We performed our analyses on cerebral cortex and hippocampus, which have been extensively studied and reported on in AD studies ^40,41^. We first looked at the overall similarity of the datasets from three time points. Interestingly, according to cluster analysis the 9 months sample strongly diverges from the other two time points in both cerebral cortex and hippocampus. Most NSPs in the two 9 months clusters show a trend in changes that is the opposite of those at 2 and 5 months. Compared with age matched controls, the majority of proteins enriched in cluster 3 in both cerebral cortex and hippocampus had increased synthesis at 2 and 5 months but decreased synthesis at 9 months in hAPP/PS1 AD mice, while most of the proteins in cluster 6 in both of the two brain regions were changed in the opposite direction (**Supplementary Figure 7&8**), suggesting distinct regulatory patterns between early (2 and 5 month) and late (9 month) stages in this AD model. RAB3C, which plays an important role in synaptic vesicle exocytosis and neurotransmitter release^42^, was enriched in cluster3 of cerebral cortex, along with Ribosomal protein S3A (RPS3A), which is associated with late-onset AD ^43^. Vesicle-trafficking protein sec22b (SEC22B) has been implicated in autophagy and its knockdown could impair autophagic flux ^44^. The synthesis of RAB3C, RPS3A and SEC22B was increased at 2 and 5 months but downregulated at 9 months in hAPP/PS1 AD mice, suggesting a positive response of selected pathways at early stages and impairment with Aβ accumulation. Overall, based on the protein synthesis profile, it appears there may an early protective phase in AD prior to severe Aβ deposition.

**Figure 7.**
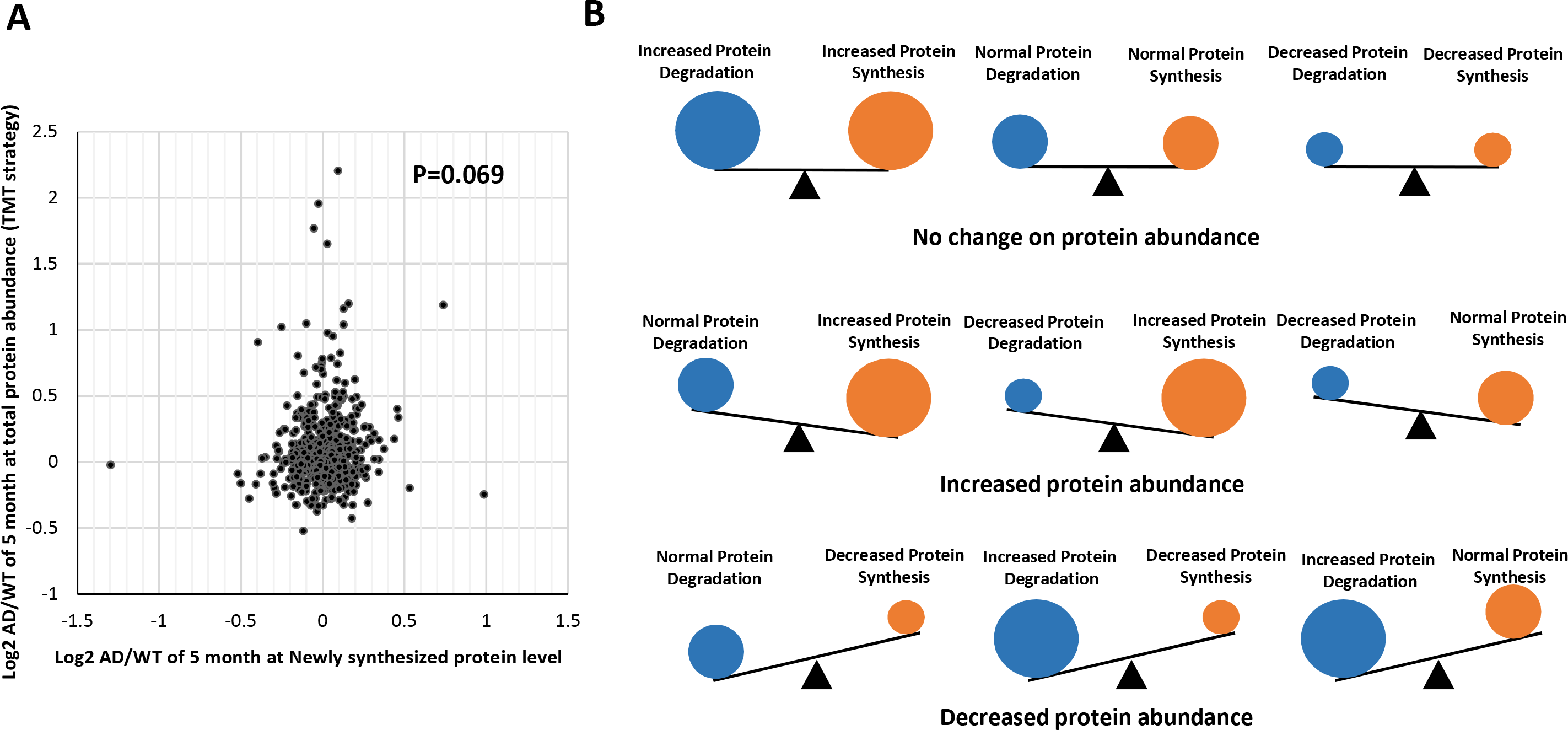
Comparison of protein synthesis and total abundance changes. **A**: Scatter plot of individual protein synthesis ratios versus abundance ratios in APP/PS1 AD mouse model versus age matched normal control. **B**: The independence between changes in protein synthesis and abundance is consistent with the notion that both synthesis and degradation contribute to the overall protein pool size.

We further assessed the significant changes in each dataset to get more information about AD pathomechanism. The overlap of differentially expressed NSPs between cerebral cortex and hippocampus is far less than that of total quantified NSPs (**Supplementary Figure 9**), suggesting that AD pathogenesis has different regulatory mechanisms in cerebral cortex and hippocampus, which is consistent with previous findings^41^. Our data also showed that almost all differentially expressed NSPs were unique for each timepoint. The pattern of differentially expressed NSPs was shown to be age-dependent and associated with the progress of AD pathogenesis.

The most interesting part of pathway analysis is that one of the significant biological processes altered at all timepoints in both brain regions is vesicle transportation. We performed network analysis on all significantly changed NSPs enriched in vesicle transportation at all timepoints and found an intricate interaction map. Multiple functional clusters had been formed by proteins associated with coat protein complex I (COPI) and mitochondria (**Supplementary Figure 10**). COPI is the main machinery in early steps of Golgi-to-endoplasmic reticulum retrograde transport and regulates APP trafficking, maturation and production of Aβ^45^. It has been reported that the moderation of COPI-dependent trafficking *in vivo* leads to a significant decrease in amyloid plaques in the cortex and hippocampus^46^. The synthesis rates of eight identified COPI subunits (COPG2, CAPZB, COPE, ARF3, ARFGAP2, RAB1A, DNC1H1 and NSF) were significantly altered in AD mice. Interestingly, they were distributed evenly over the three different timepoints. Mitochondria is an important source of fuel for many biological processes including vesicle transportation. We have identified that seven subunits (ATP5O, ATP5A, COX5A, COX4I1, MTCO2, SUCLA2 and UQCRC2) of the electron transport chain were significantly regulated in the vesicle transport biological process of AD. Among them, the downregulated subunits were all seen at 9 months, indicating the declined activity of respiratory chain and energy production at late stage AD. Our analysis further supports the conclusion that different proteins in the same functional groups are regulated with the progress of AD. These findings support previously reported findings that COPI-dependent trafficking as well as the activity of mitochondria are important factors in AD, and they merit further study^46^.

Our findings are consistent with the emerging hypothesis that endosome function is altered in Alzheimer’s disease ^47^. For example, it was observed that the volumes of endosomes in patients with AD are up to 32-fold larger than normal, implying a marked increase in endocytic activity^48^. Similarly, it has been shown that the alterations in sortilin-related receptor L (SORL), bridging integrator 1 (BIN1) and CD2 associated protein (CD2AP) all cause endosomal enlargement^49^. Our study contributes to this idea by demonstrating that proteins related to endosomal metabolism were abnormally regulated prior to Aβ accumulation and symptoms. Additionally, different proteins involved in the same steps of endosomal metabolism were regulated with AD progress.

It has been reported that changes in protein turnover and protein abundance are not consistent^50^. In our study, a measurable change in newly synthesized proteins may indicate molecular scenarios that differ from the measurement of total protein abundance. To confirm this hypothesis, we compared our NSPs dataset with a previously published study, which quantified protein abundance based on tandem mass tag (TMT) proteomic analysis in the same AD mouse model^22^. This comparison allows us to further explore the relationship between the kinetics of protein synthesis and protein abundance. We found only a modest correlation between changes in protein synthesis and total proteins in cerebral cortex of 5months APP/PS1 AD mice (Figure 7A, P=0.069), indicating that many proteins with increased synthesis did not in fact increase the total protein abundance, and vice versa. The possible reason might be that protein degradation is altered in AD, since both protein synthesis and degradation contribute to protein pool size (Figure 7B). Our analysis indicated that the changes in protein synthesis and protein abundance in a system are largely independent parameters. Since NSPs are, in theory, the first to respond to pertubations, we believe that the measurement of protein synthesis is a more direct way to monitor regulation changes. The novel integration of NSPs and protein abundance proteomic datasets could be applied to all animal models of disease to provide a greater understanding of homeostasis in human diseases.

In summary, this study presents a direct, sensitive, accurate and versatile workflow for the quantitation of newly synthesized proteins with individual protein resolution in mouse disease model. This method is compatible with large-scale inquiries and allows protein synthesis to be measured over a narrow window of time by using *in vivo* HAHA/AHA labeling through diet. Using this strategy, we identified a comprehensive dataset of NSPs at 2, 5 and 9 months of APP/PS1 AD mice and demonstrated that patterns of differentially expressed NSPs are age-dependent. Our results point to the dysfunction of vesicle transport prior to Aβ accumulation and symptoms, which may progressively impact the entire protein network and thereby drive neurodegeneration. Taken together, our findings present a potential pathomechanism by which Aβ accumulation interferes with cellular functions through dysregulation of vesicle transportation, which provides unique insights into the mechanism that underlie proteomic changes in AD.

## ACKNOWLEDGEMENTS

We thank Dr. C. Delahunty for critical reading. This work was supported by the National Institute of Health grants R03AG047957 and P41 GM103533.

