## supplementary figures 1-10 for "A Temporal Quantitative Profiling of Newly Synthesized Proteins during Aβ Accumulation"

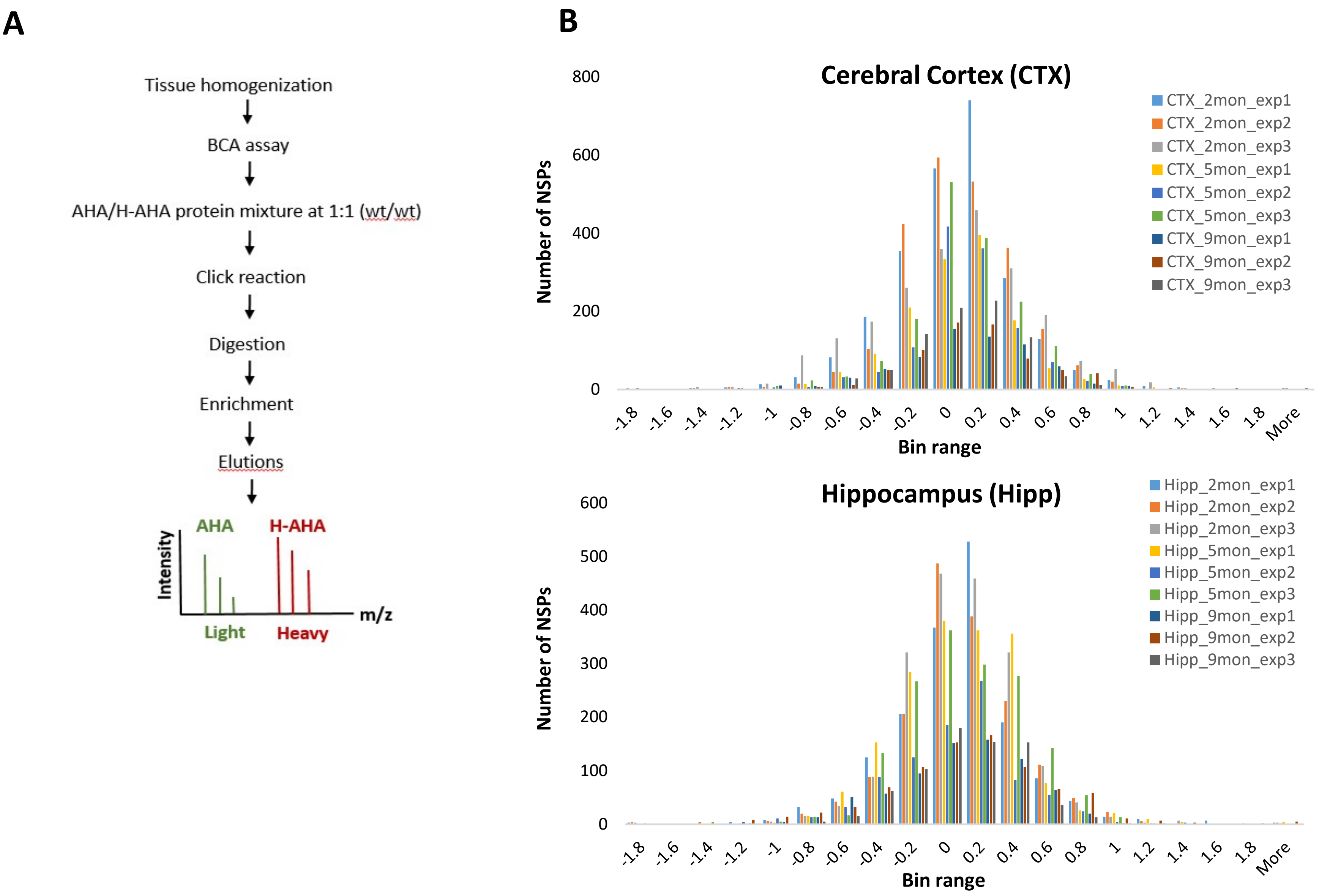


***Supplementary Figure 1***: Workflow of sample preparation for MS analysis. Click reaction was performed on the proteins to covalently add a biotinalkyne to the AHA molecule incorporated in the proteins. Tryptic digestion was performed. Peptides with the AHAbiotin-alkyne modification were enriched with neutravidin beads. The beads were washed to remove unmodified peptides. The modified peptides were eluted off the beads and the eluate was analyzed by MS to identify newly synthesized proteins. **B.** Normal distribution of replicates of hippocampus and cerebral cortex from different time points.

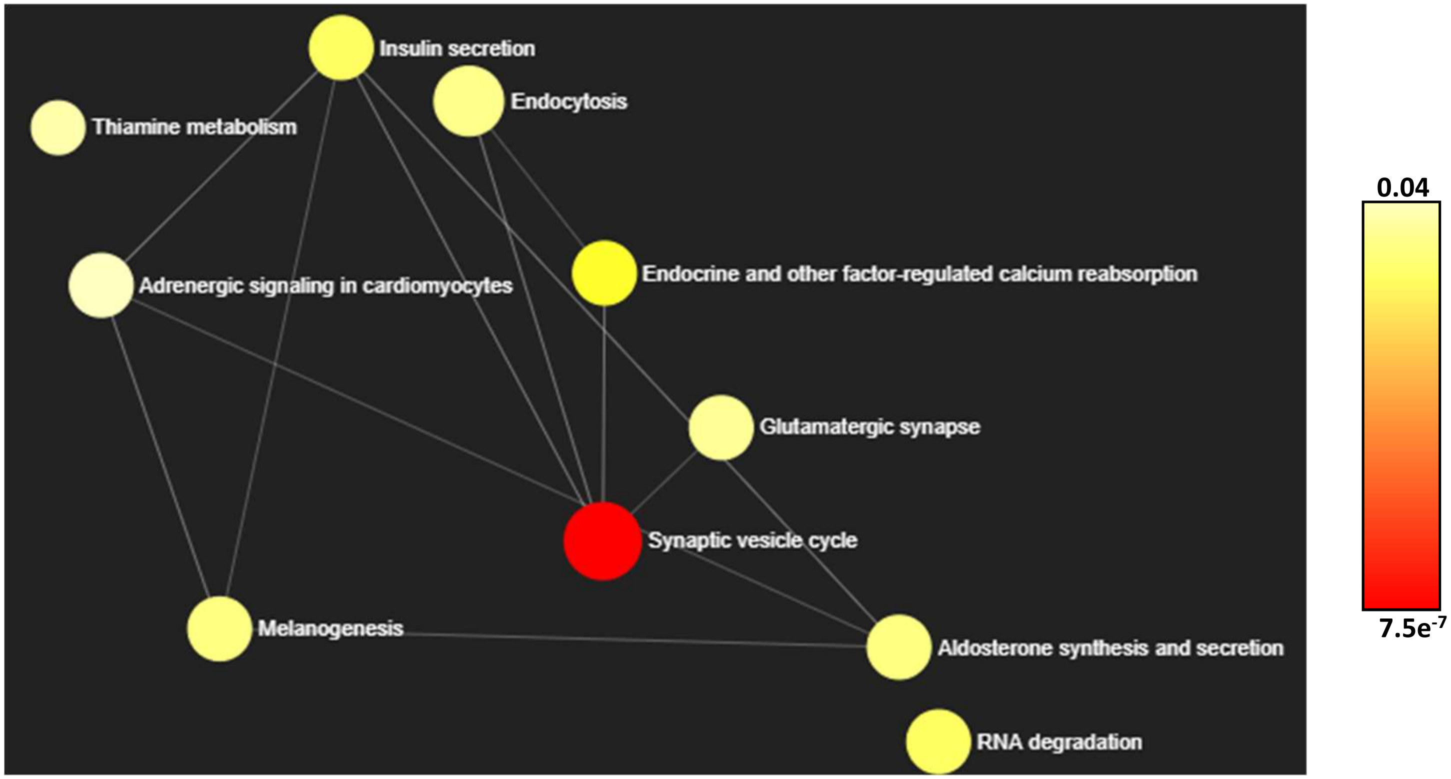


***Supplementary Figure 2***. KEGG pathway analysis on cluster 3 (see supplementary table 3) of cerebral cortex. P value of each enriched pathways are represented by gradient colors. All analysis here was done by NetworkAnalyst 3.0 ([https://www.networkanalyst.ca/](https://www.networkanalyst.ca/)).

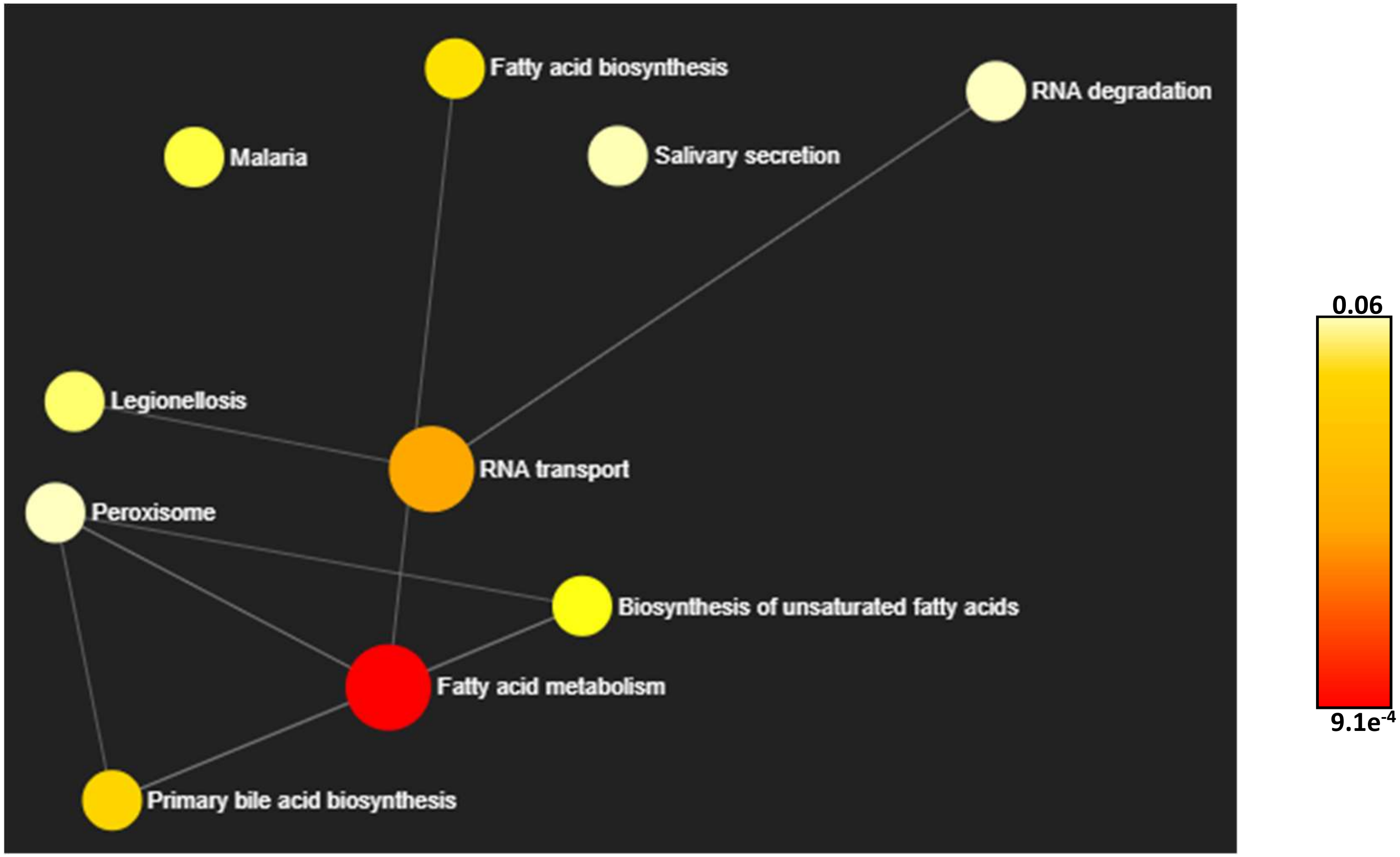


***Supplementary Figure 3***. KEGG pathway analysis on cluster 6 (see supplementary table 3) of cerebral cortex. P value of each enriched pathways are represented by gradient colors. All analysis here was done by NetworkAnalyst 3.0 (https://www.networkanalyst.ca/).

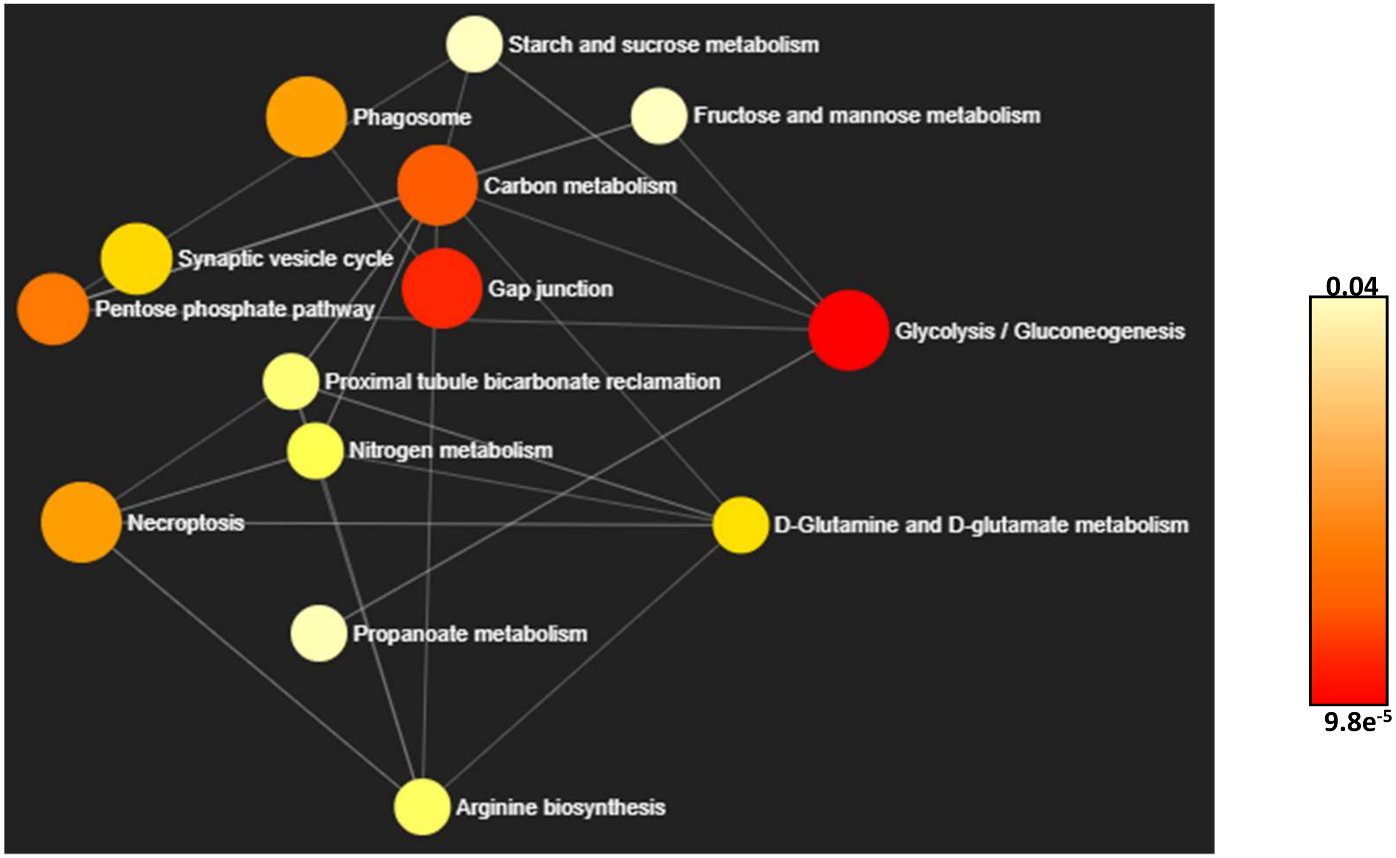


***Supplementary Figure 3***. KEGG pathway analysis on cluster 3 (see supplementary table 4) of hippocampus. P value of each enriched pathways are represented by gradient colors. All analysis here was done by NetworkAnalyst 3.0 (https://www.networkanalyst.ca/).

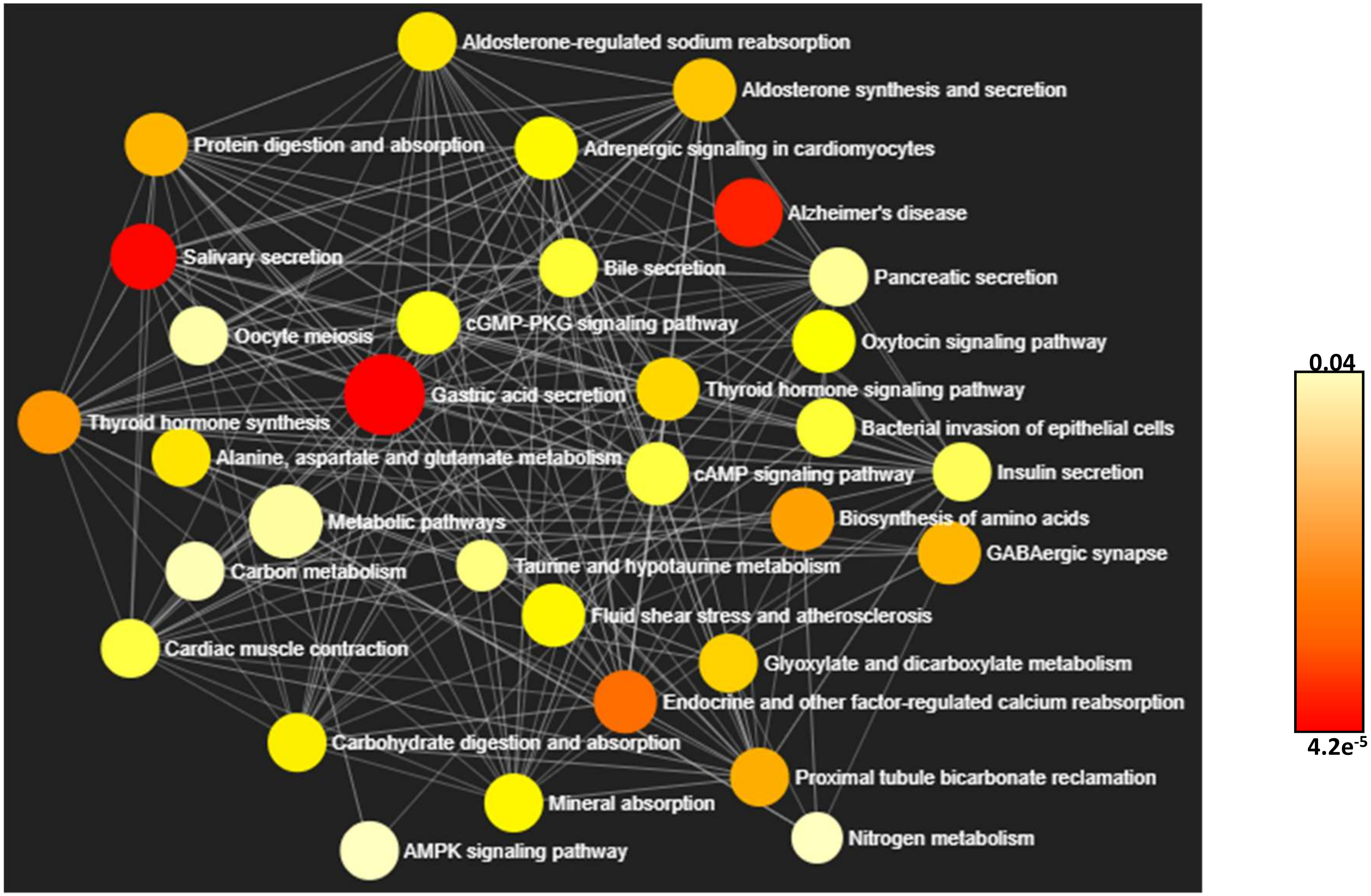


***Supplementary Figure 5.*** KEGG pathway analysis on cluster 6 (see supplementary table 4) of hippocampus. P value of each enriched pathways are represented by gradient colors. All analysis here was done by NetworkAnalyst 3.0 (https://www.networkanalyst.ca/).

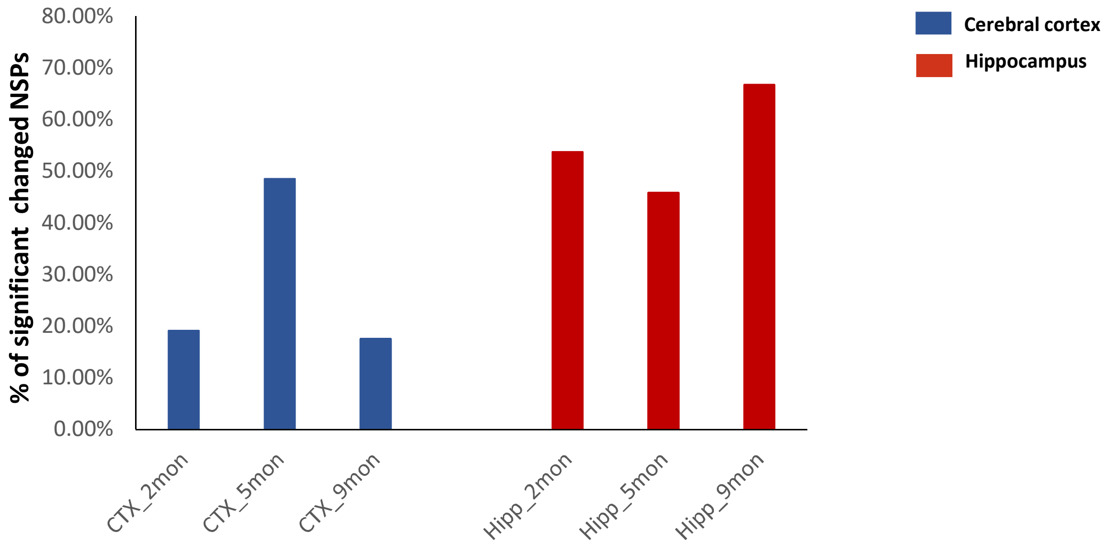


***Supplementary Figure 6.*** The percentage of significantly changed NSPs enriched in vesicle mediated transport/membrane bounded vesicle.

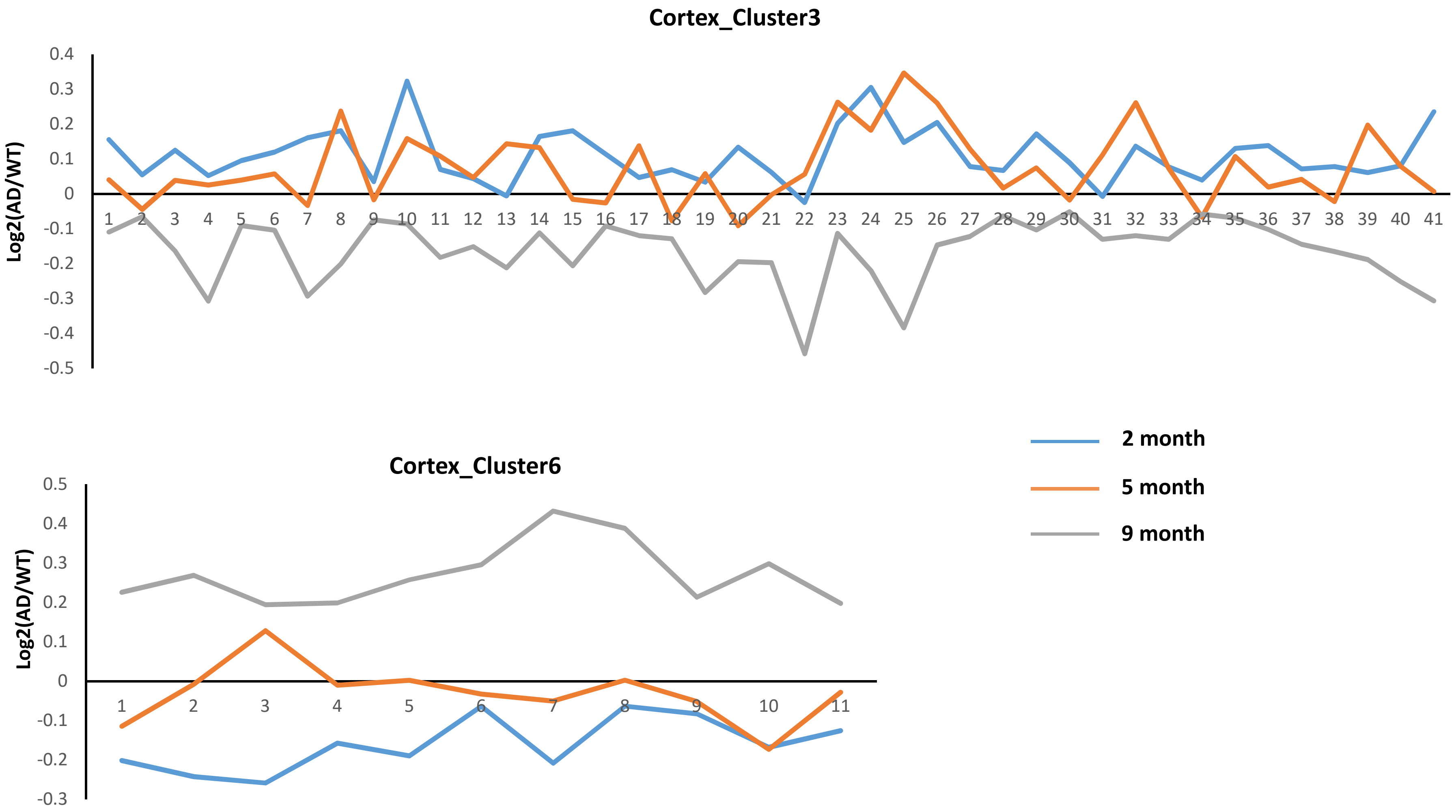


***Supplementary Figure 7.*** Ploting of the data points from cluster 3 and 6 of cerebral cortex.

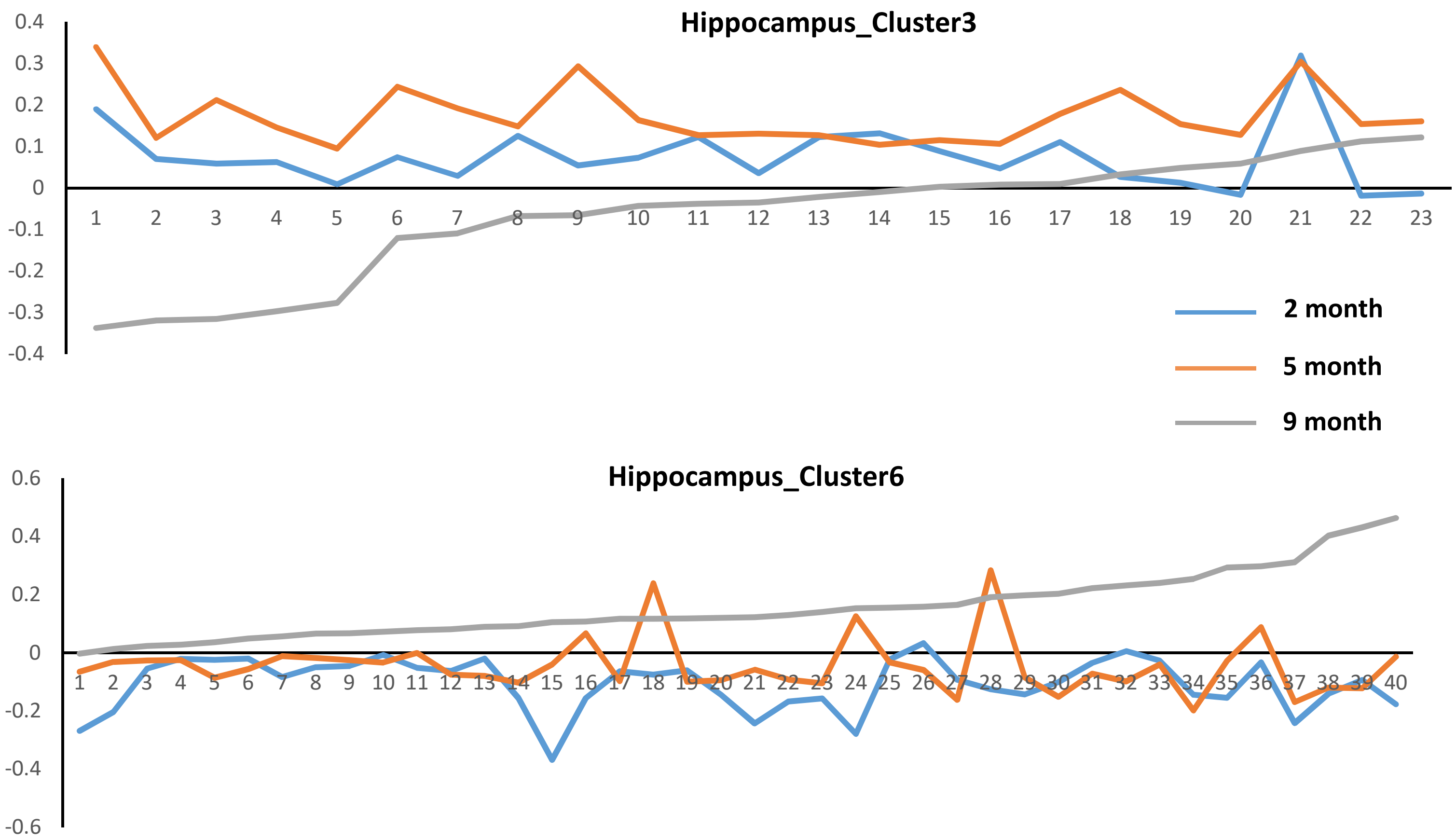


***Supplementary Figure 8.*** Ploting of the data points from cluster 3 and 6 of hippocampus

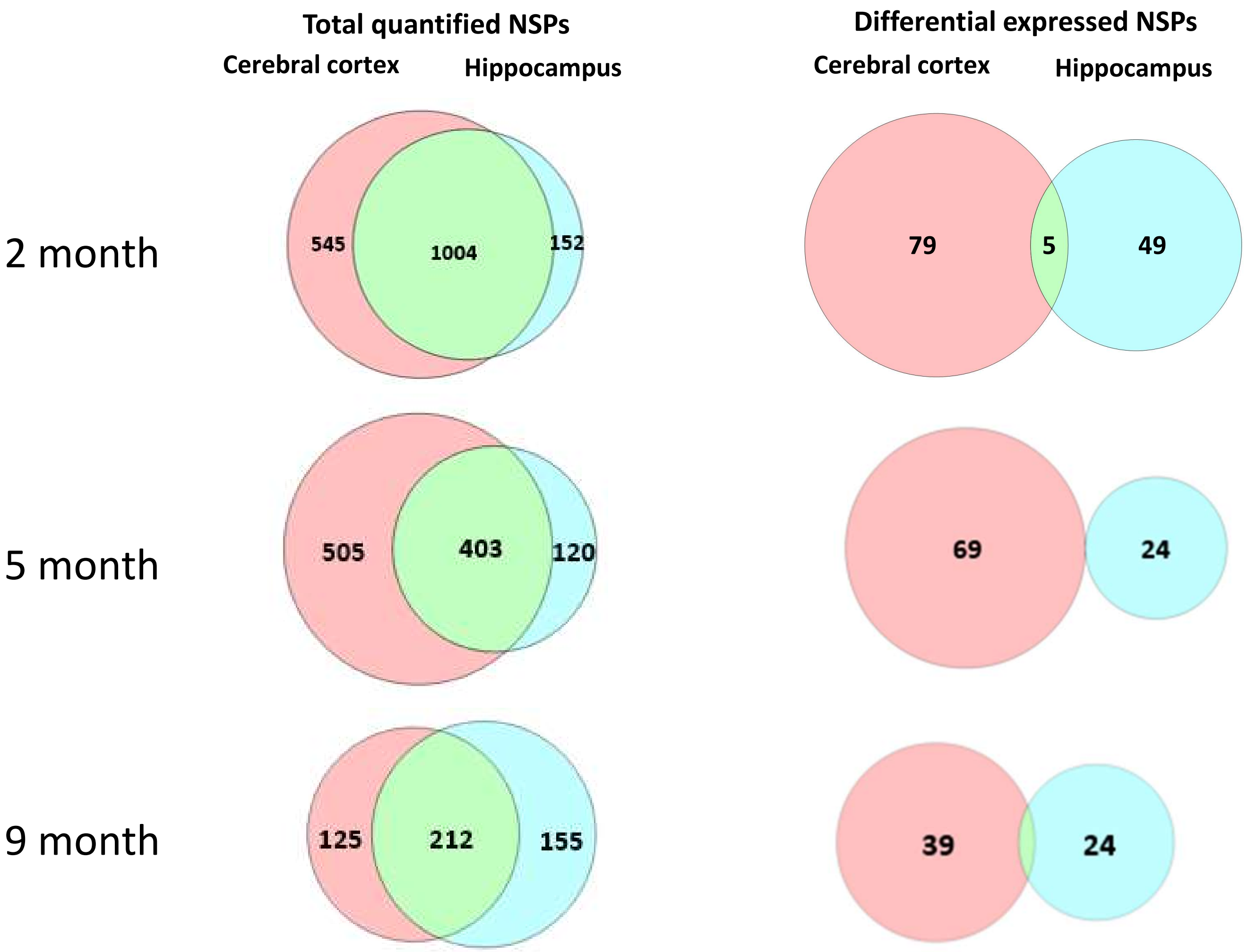


***Supplementary Figure 9.*** Venn diagram of total quantified NSPs and differential expressed NSPs for each time point.

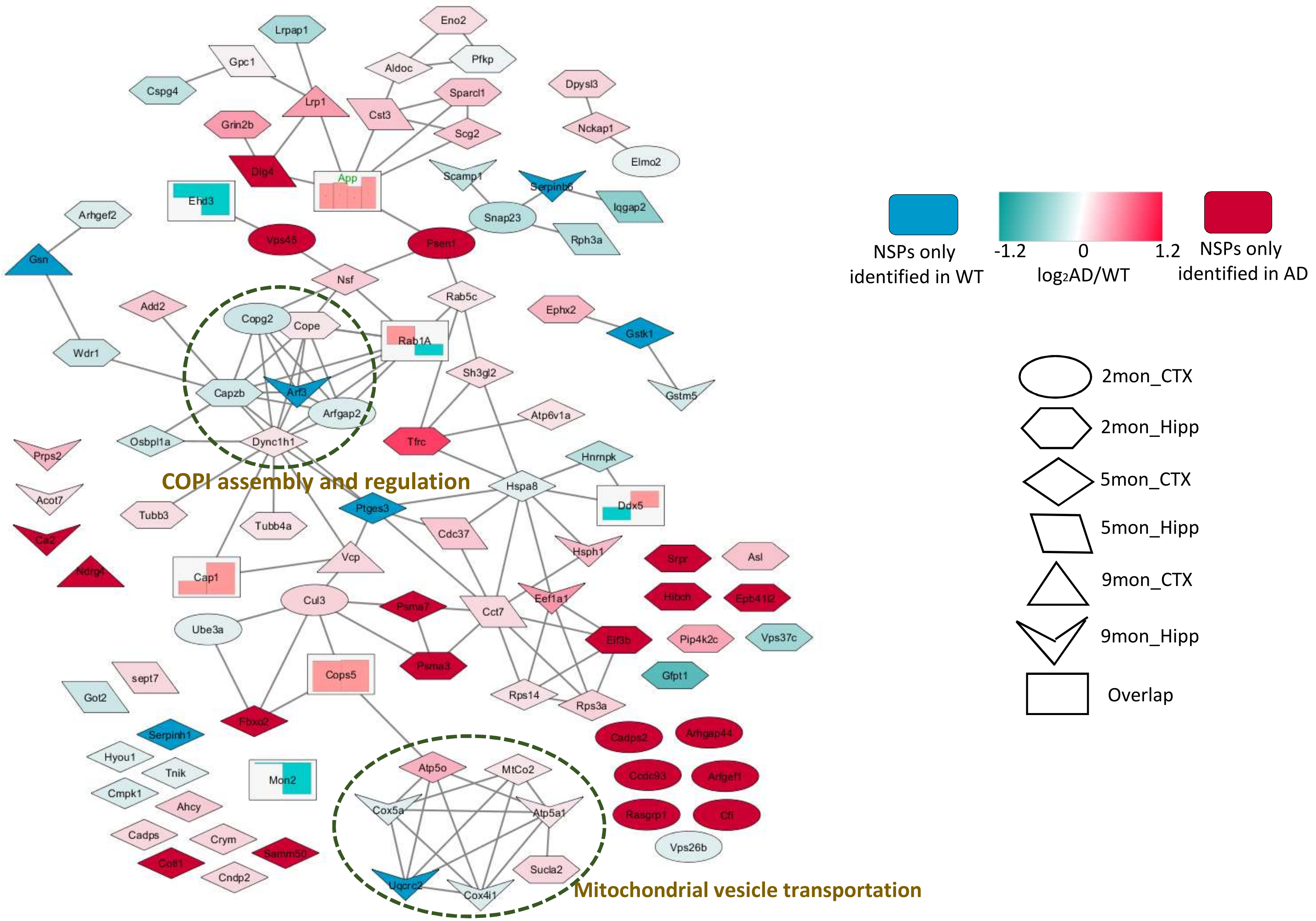


***Figure 10***. STRING network analysis of significant altered NSPs, which were enriched in vesicle related processes. Only interactions with a STRING score ≥ 0.7 are shown. Node color are linearly related to fold-change (Rectangular node excluded). Node shape represents different timepoints and brain region. The shape "rectangular " represents the overlapped NSPs fold-changes indicated in bar graph. The column in red means increased synthesis while that in green means down regulation. The overlapped NSPs were listed here with bar graph annotated from left to right. **APP**: 2mon_CTX; 2mon_Hipp; 5mon_Hipp; 9mon_CTX; **Cap1**: 2mon_Hipp, 9mon_CTX; **Cops5**: 2mon_Hipp, 5mon_Hipp; **Ddx5**: 5mon_CTX; 9mon_Hipp; **Ehd3**: 5mon_CTX; 9mon_Hipp; **Mon2**: 2mon_Hipp; 5mon_CTX; **Rab1A**: 9mon_CTX; 9mon_Hipp
