## supplementary table 1 for "A Temporal Quantitative Profiling of Newly Synthesized Proteins during Aβ Accumulation"

***Supplementary Table 1***:Identification results

| Timepoint | Replicate | Content | Cortex | | Hippocampus | |
|---|---|---|---|---|---|---|
| | | | Protein | peptide | Protein | peptide |
| 2mon | 1 | H-AD/L-WT | 2,678 | 19,904 | 1,852 | 11,074 |
| | 2 | L-AD/H-WT | 2,548 | 15,798 | 1,841 | 10,963 |
| | 3 | L-AD/H-WT | 2,520 | 17,934 | 2,127 | 12,884 |
| 5mon | 1 | H-AD/L-WT | 1,509 | 7,971 | 2,183 | 12,174 |
| | 2 | L-AD/H-WT | 1,349 | 8,746 | 1,034 | 4,889 |
| | 3 | L-AD/H-WT | 1,815 | 12,883 | 2,297 | 12,218 |
| 9mon | 1 | L-AD/H-WT | 880 | 2,832 | 869 | 2,965 |
| | 2 | H-AD/L-WT | 755 | 3,140 | 1,002 | 3,780 |
| | 3 | H-AD/L-WT | 961 | 4,856 | 810 | 2,917 |
