## supplementary table 2 for "A Temporal Quantitative Profiling of Newly Synthesized Proteins during Aβ Accumulation"

***Supplementary Table 2***: Quantification efficiency. Only proteins quantified with sigma less than or equal 0.5 were counted in this statistics.

| Timepoint | Replicate | Content | Cortex | Hippocampus |
|---|---|---|---|---|
| 2mon | 1 | H-AD/L-WT | 92.9% | 90.6% |
| | 2 | L-AD/H-WT | 91.8% | 90.8% |
| | 3 | L-AD/H-WT | 85.3% | 89.0% |
| 5mon | 1 | H-AD/L-WT | 90.9% | 85.7% |
| | 2 | L-AD/H-WT | 92.1% | 86.8% |
| | 3 | L-AD/H-WT | 90.2% | 89.3% |
| 9mon | 1 | L-AD/H-WT | 87.2% | 85.4% |
| | 2 | H-AD/L-WT | 91.3% | 85.9% |
| | 3 | H-AD/L-WT | 88.6% | 89.8% |
